# Loss of Muscleblind Splicing Factor Shortens *C. elegans* Lifespan by Reducing the Activity of p38 MAPK/PMK-1 and Transcription Factors ATF-7 and Nrf/SKN-1

**DOI:** 10.1101/2021.02.22.432374

**Authors:** Olli Matilainen, Ana R. S. Ribeiro, Jens Verbeeren, Murat Cetinbas, Ruslan I. Sadreyev, Susana M. D. A. Garcia

## Abstract

Muscleblind-like splicing regulators (MBNLs) are alternative splicing factors that have an important role in developmental processes. Dysfunction of these factors is a key contributor of different neuromuscular degenerative disorders, including Myotonic Dystrophy type 1 (DM1). Since DM1 is a multisystemic disease characterized by symptoms resembling accelerated aging, we asked whether MBNLs regulate cellular processes required to maintain normal lifespan. By utilizing the model organism *Caenorhabditis elegans*, we found that loss of MBL-1 (the sole ortholog of mammalian MBNLs), which is known to be required for normal lifespan, shortens lifespan by decreasing the activity of p38 MAPK/PMK-1 as well as the function of transcription factors ATF-7 and SKN-1. Furthermore, we show that mitochondrial stress caused by knockdown of mitochondrial electron transport chain components promotes the longevity of *mbl-1* mutants in a partially PMK-1-dependent manner. Together, the data establish a mechanism of how DM1-associated loss of muscleblind affects lifespan. Furthermore, this study suggests that mitochondrial stress could alleviate symptoms caused by the dysfunction of muscleblind splicing factor, creating a potential approach to investigate for therapy.

**Reviewer token for the RNA-seq data (GEO: GSE146801):** wvataksittaffcj

## Introduction

The human Muscleblind-like (MBNL) protein family consists of MBNL1, MBNL2 and MBNL3. These evolutionarily conserved splicing factors have an important role during development as they regulate the alternative exon choice during terminal muscle differentiation (PASCUAL *et al*. 2006). Due to their role as regulators of cellular alternative splicing programs, dysfunction of MBNLs has been found to be implicated in many degenerative diseases such as huntington’s disease-like 2, spinocerebellar ataxia type 8 and polyglutamine disorders (FERNANDEZ-COSTA *et al*. 2011). However, MBNL dysfunction is best known for its role in myotonic dystrophies.

Myotonic dystrophy type 1 (DM1) and type 2 (DM2) are multisystemic diseases caused by 50-4.000 CUG trinucleotide repeat expansions in 3′ untranslated region (UTR) of the myotonic dystrophy protein kinase (*DMPK*) gene and 55-11.000 CCUG repeats in the first intron of the cellular nucleic acid-binding protein (*CNBP*) gene, respectively. DM1 is the most prevalent muscular dystrophy in adults, and its clinical features are more severe compared to DM2 (MEOLA AND CARDANI 2015). At the molecular level, repeat-containing RNAs form imperfect double-stranded structures that sequester, and thereby disrupt the function of MBNLs and several other RNA binding factors (MILLER *et al*. 2000; FARDAEI *et al*. 2001), which leads to splicing defects (RANUM AND COOPER 2006). Importantly, *Mbnl1* and *Mbnl2* knockout mice develop RNA splicing abnormalities that are characteristic of DM1 (KANADIA *et al*. 2003; CHARIZANIS *et al*. 2012; SUENAGA *et al*. 2012), demonstrating that the loss of MBNL splicing factors is a central contributor of DM1 pathogenesis. Since DM1 manifests with phenotypes commonly associated with aging (e.g. cataracts, muscular weakness, atrophy, insulin resistance and metabolic dysfunction), it has also been considered as a disorder of accelerated aging (MATEOS-AIERDI *et al*. 2015). Thus, the function of MBNL splicing factors may play an important regulatory role in aging.

MBNLs are well conserved factors across animals. In *Caenorhabditis elegans,* MBL-1, the sole ortholog of mammalian MBNL proteins binds CUG and CCUG repeats (SASAGAWA *et al*. 2009; GARCIA *et al*. 2014), and is required for normal mRNA splicing (NORRIS *et al*. 2017; THOMPSON *et al*. 2019). MBL-1 appears to be required for normal muscle structure and function as well as for proper formation of neuromuscular junction synapses and dendrite morphogenesis (WANG *et al*. 2008; SPILKER *et al*. 2012; ANTONACCI *et al*. 2015), indicating that the loss of MBL-1 recapitulates DM1 patient phenotypes in *C. elegans*. Notably, it has been shown that the loss of MBL-1 shortens *C. elegans* lifespan (WANG *et al*. 2008; SASAGAWA *et al*. 2009), but the mechanism behind this phenotype is not known.

Here we used *C. elegans* to elucidate the mechanism of how loss of MBL-1 shortens lifespan. We found that MBL-1-deficient animals have reduced activity of the conserved p38 mitogen-activated protein kinase (MAPK) (PMK-1 in *C. elegans*) and its downstream transcription factors ATF-7 and SKN-1, which results in shortened lifespan. Furthermore, we demonstrate that mitochondrial stress activates a prolongevity response in *mbl-1* mutants, which is partially p38 MAPK/PMK-1-dependent. These findings establish PMK-1, ATF-7 and SKN-1 as mechanistic links between the developmentally regulated muscleblind splicing factor and lifespan, and establish mitochondria as potential therapeutic targets to treat aging-associated symptoms of DM1.

## Materials and methods

### *C. elegans* strains and maintenance

For all experiments *C. elegans* were maintained at 20°C on NGM plates (peptone, P4963, Merck; agar, A4550, Merck; NaCl, 746398, Merck). The N2 (Bristol) strain was used as the wild-type. N2, *tir-1(qd4)* (ZD101), *pmk-1(km25)* (KU25), *gst-4p::gfp* (CL2166) were obtained from the Caenorhabditis Genetics Center (CGC). *mbl-1(tm1563)* strain was received from National BioResource Project (NBRP), and outcrossed five times with N2. *mbl-1(tm1563);gst-4p::gfp* (GAR132) and *pmk-1(km25);mbl-1(tm1563)* (GAR139) crosses as well as extrachromosomal *mbl-1* overexpressing animals (GAR152: (iceEx51[*mbl-1p(1)::mbl-1-isoA::mCherry*] [*mbl-1p(2)::mbl-1-isoA::mCherry*] [*mbl-1p(1)::mbl-1-isoB::mCherry*] [*mbl-1p(2)::mbl-1-isoB::mCherry*] [*mbl-1p(1)::mbl-1-isoC::mCherry*] [*mbl-1p(2)::mbl-1-isoC::mCherry*]) were created within this study.

### Pseudomonas aeruginosa assay

3.5 cm NG plates were seeded with 40 µL of overnight grown *Pseudomonas aeruginosa* (PA14) suspension and incubated at 37°C for 24 h (TAN *et al*. 1999). 20 µL of SDS 2% was added to the edges of the plate to prevent the escape of the animals. *C. elegans* were transferred to PA14 plates at L4 stage and incubated at 25°C. Animals were scored daily for survival based on their ability to respond to touch. Animals dehydrated on the wall of the plate were censored from the analysis. For Western blot upon PA14 treatment, Animals were transferred to PA14 plates at L4 stage and collected 24 hours later as day 1 adults.

### RNA interference (RNAi)

*tir-1*, *pmk-1*, *atf-7*, *skn-1* and *cco-1* RNAi clones were taken from either Ahringer or Vidal RNAi library. *daf-16* (#34833) and *daf-2* RNAi (#34834) clones were obtained from Addgene. *mbl-1* RNAi was cloned from *C. elegans* cDNA by using 5’-CTTGAGCTCTTCGACGAAAACAGTAATGCCGC-3’ and 5’-CTTCTCGAGCTAGAATGGTGGTGGCTGCATG-3’ primers. RNAi was performed using the feeding protocol as described earlier (TIMMONS *et al*. 2001). Lifespan experiments were initiated by letting gravid hermaphrodites (P0 generation) to lay eggs on RNAi/NGM plates, and P1 generation was scored for lifespan. In some lifespan experiments animals were bleached and let to hatch overnight in M9 before plating P1 generation as L1 larvae to experimental plates. These two alternative ways to initiate lifespan did not affect the conclusions made from the experiments. Since *mbl-1* is expressed mainly in neurons, P0s were put to *mbl-1* RNAi at L4 stage in order to maximize RNAi efficiency in P1 generation, which was scored for lifespan. For qRT-PCR and Western blot animals were bleached and let to hatch overnight in M9 before plating L1 larvae for experimental plates.

### Generation of *mbl-1* overexpression strains

GAR152 was generated by microinjection of six different transgenic construct at 15 ng/μl concentration each. These transgenes were expressed as extrachromosomal arrays and express three different isoforms (K02H8.1a; isoform A, K02H8.1b; isoform B, and K02H8.1c, isoform C) fused to mCherry, under two different endogenous promoters; *mbl-1p(1)* and *mbl-1p(2)*. Constructs were made through modification of a plasmid expressing *mbl-1* isoform A fused to mCherry under the *unc-54* promoter (GARCIA *et al*. 2014). PCR-amplified fragments from wild-type *C. elegans* genome, representing two endogenous *mbl-1* promoter regions, replaced the *unc-54* promoter. For promoter 1, *mbl-1p(1)*, primers 5’-TACGCATGCAGGCCCTATATATTCCATCTCAAT-3’, containing a *Sph*I site, and 5’-TACGGATCCTCTGAAAAGTAGGAAAAAGATTGGC, containing a *Bam*HI site were used. For promoter two, *mbl-1p(2)*, primers 5′-AACTGCAGGTGCAATGGGCTACTGATCTCC-3′ and 5′-CGGGATCCCATTCCGTCACTTGCAAAGAAC-3′ were used, containing *Pst*I and *Bam*HI sites, respectively. Plasmids expressing isoforms B and C were derived from these constructs through PCR using forward primers 5′-CAGCTACAAACTGCCGC CT-3′ (isoform B) and 5′-GGAGCTGTACCAATGAAGCGAC-3′ (isoform C), and reverse primer 5′-CTGATTCACTGCCGCTGCTGTATAAG-3′.

### Western blot

Animals were collected at indicated age and frozen in liquid nitrogen. Animals were lysed in protease inhibitor cocktail (abcam, #ab65621) and 3 x phosphatase inhibitor (Thermo Scientific, #78420)-supplemented RIPA buffer (bioWORLD) with micropestle in 1.5 ml Eppendorf tubes. Lysates were resolved on 4–15% precast polyacrylamide gels (Bio-Rad). Phospho-p38 MAPK (used with 1:1000 dilution) was purchased from Cell Signaling Technology (#4511S) and α-tubulin antibody (used with 1:5000 dilution) from Merck (T5168). Western blots were quantified by using Fiji (see S6 Table for Western blot quantifications).

### RNA sequencing

N2 and *mbl-1(tm1563)* strains were synchronized by bleaching and plated as L1 larvae on RNAi plates seeded with HT115 bacteria carrying empty vector (control vector for RNAi). To prevent the hatching of the progeny, animals were transferred to plates containing 10 μM of 5-fluorouracil (Sigma) at the L4 stage. Animals were collected at day 2 of adulthood (three biological replicates for both strains) and frozen in liquid nitrogen. Total RNA was extracted with TRIzol Reagent (Ambion) and assessed for degradation using Agilent 2100 Bioanalyzer. Illumina Truseq stranded polyA-mRNA library was prepared and sequenced for 86 cycles at the DNA Sequencing and Genomics laboratory (Institute of Biotechnology, University of Helsinki). The 6 samples were multiplexed and sequenced on one lane of Illumina NextSeq 500, yielding circa 18–20 million reads per sample. Sequencing reads were mapped to the *Caenorhabditis elegans* reference transcriptome (WBcel235 assembly) using STAR (DOBIN *et al*. 2013). Read counts over transcripts were calculated using HTSeq (ANDERS *et al*. 2015). For differential expression analysis we used the EdgeR method (ROBINSON *et al*. 2010) and classified genes as differentially expressed based on the cutoffs of 2-fold change in expression value and false discovery rates (FDR) below 0.05.

### Lifespan analysis

*C. elegans* lifespan experiments were done at 20°C. At L4 larval stage animals were transferred to plates containing 10 µM of 5-Fluorouracil (Sigma) in order to prevent progeny production. Animals that had exploded vulva or that crawled off the plate were censored. Animals were counted as dead if there was no movement after poking with a platinum wire. Lifespans were checked every 1-3 days. For lifespan data, mean lifespan ± standard error (s.e.) is reported (see S1-S2 Tables).

### Quantitative RT-PCR (qRT-PCR)

Animals were collected at indicated age and frozen in liquid nitrogen. TRIzol Reagent (Ambion) was used to extract RNA. cDNA synthesis was done with QuantiTect Reverse Transcription Kit (Qiagen) and qRT-PCR reactions were run with the SYBR Green reagent (Roche) using Lightcycler 480 (Roche). qRT-PCR data was normalized to the expression of *cdc-42* and *pmp-3*. qRT-PCR oligos used in this study are provided in S8 Table. qRT-PCR experiments were performed with three biological replicates (see S7 Table for raw qRT-PCR data), and repeated at least twice with similar results. Statistical significances were analyzed by using Student’s t-test or two-way ANOVA.

### Splicing assay

Animals were collected at indicated age and frozen in liquid nitrogen. TRIzol Reagent (Ambion) was used to extract RNA. cDNA synthesis was done with QuantiTect Reverse Transcription Kit (Qiagen). Splicing of *unc-43* and *unc-104* mRNAs was analyzed by running PCR products in agarose gels. Oligos used for PCR are provided in S9 Table. Splicing assays were performed with three biological replicates. Agarose gels were quantified by using Fiji. Statistical significances were analyzed by using two-way ANOVA.

### Fluorescent imaging and oxidative stress assay

*gst-4p::GFP* reporter strains (wild-type and *mbl-1(tm1563)* background) were grown on EV RNAi plates and imaged as day 1 adults with a standard stereomicroscope with fluorescent light source (Zeiss). For the oxidative stress, 5 % H2O2 (in H2O, H2O used as a control) was pipetted to EV plates containing 1 day adult *gst-4p::gfp* reporter animals. Worms were washed after 20 minutes, and let to recover on EV plates for 5 hours before imaging. GFP fluorescence was quantified by using Fiji (see S7 Table for GFP quantifications).

### Statistical analysis

Statistical analyses for qRT-PCR data was carried out in GraphPad Prism or Excel, and the data represents the mean of 3 biological replicates ± SD. Statistical analyses for Western blot data, splicing assays (agarose gels) and GFP fluorescence were carried out in GraphPad Prism. Statistical details can be found in the figures and figure legends. Statistical analyses for lifespan experiments were carried out in R by using the Cox-proportional hazard regression. Statistical details for the lifespan data can be found in S1 and S2 Tables.

### Data and reagent availability

Strains and plasmids are available upon request. The Gene Expression Omnibus (GEO) accession number for the RNA-seq data originating from this study is GSE146801.

## Results

### Loss of MBL-1 leads to shortened lifespan and reduced expression of p38 MAPK/PMK-1-regulated genes

To examine how the loss of MBL-1 affects *C. elegans* lifespan, we utilized a strain carrying the *mbl-1(tm1563)* allele (513 bp deletion) that eliminates an exon shared among all *mbl-1* isoforms and creates a putative null allele. This mutant has been shown to shorten lifespan (SASAGAWA *et al*. 2009). We repeated this experiment, performing lifespan assays on two *E. coli* strains, HT115 and OP50, which are both widely used in *C. elegans* experiments. We found that *mbl-1(tm1563)* mutant animals have significantly shorter lifespan on both bacterial strains compared to wild-type *C. elegans* (N2) (Fig 1A and S1-S2 Tables). Interestingly, loss of MBL-1 has a more drastic decrease in lifespan on HT115 compared to OP50. One potential explanation is that the microbiome could modulate lifespan partly through MBL-1. As many experiments in this study take advantage of RNAi, all following experiments (with the exception of the *Pseudomonas aeruginosa* resistance assay) were performed using HT115 bacteria. After confirming the earlier finding that *mbl-1(tm1563)* mutants have shortened lifespan (SASAGAWA *et al*. 2009) (Fig 1A and S1-S2 Tables), we asked whether knockdown of *mbl-1* produces a similar phenotype. Notably, it has been shown earlier that *mbl-1* RNAi shortens lifespan (WANG *et al*. 2008). We also found that *mbl-1* RNAi shortens N2 lifespan (Fig 1B and S1-S2 Tables), thus further demonstrating that disruptions to MBL-1 function affects lifespan. Interestingly, also simultaneous overexpression of three mCherry-tagged *mbl-1* isoforms (Fig S1A-S1B) under endogenous *mbl-1* promoters shortens lifespan compared to N2 (Fig 1C and S1-S2 Tables), indicating that an optimal level of this developmentally-regulated splicing factor is required for normal lifespan. Next, we asked whether MBL-1 activity decreases with aging. For this purpose, we scanned the *unc-43* and *unc-104* genes, known targets of MBL-1-mediated splicing (NORRIS *et al*. 2017), for age-related changes in their alternative splicing patterns (Fig S2A). PCR analysis revealed that MBL-1 is required for exon exclusion events in both *unc-43* and *unc-104* (Fig S2B-S2E, S3 Table). Day 4 adult N2 shows enrichment of longer *unc-43* isoform, which is observed with *mbl-1(tm1563)* mutants (Fig S2B-S2C, S3 Table), whereas *unc-104* splicing shows a similar enrichment already in day 2 adult N2 (Fig S2D-S2E, S3 Table). These data suggest that MBL-1 activity decreases upon aging.

**Fig. 1.**
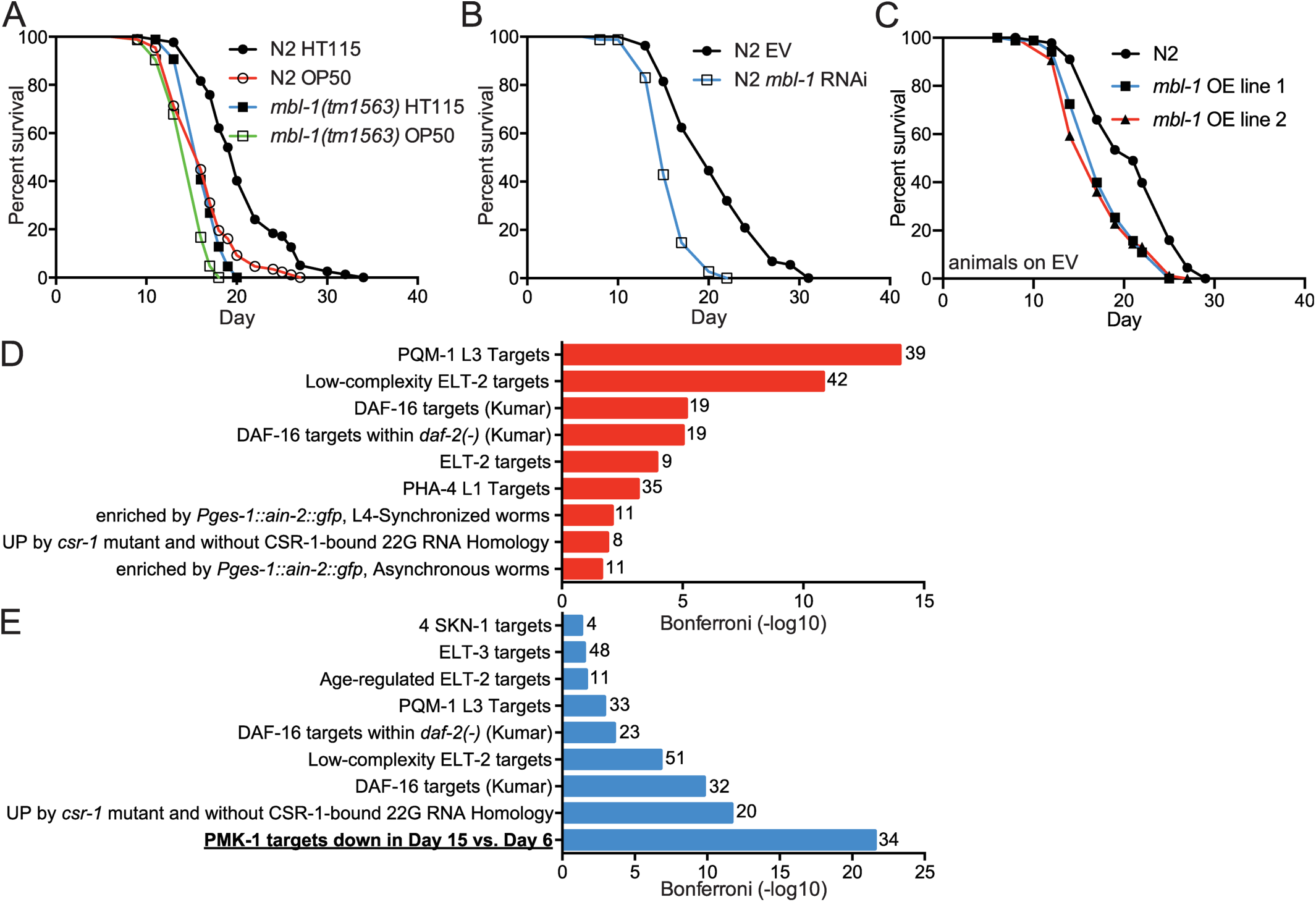
The loss of MBL-1 shortens lifespan and reduces the expression of p38 MAPK/PMK-1 target genes. (A) *mbl-1(tm1563)* mutants have shortened lifespan compared to wild-type (N2) on both HT115 (p < 0.01) and OP50 (p < 0.01) *E. coli*. (B) *mbl-1* RNAi shortens lifespan compared to empty vector (EV, control RNAi)-treated N2 (p < 0.01). (C) Strains expressing extrachromosomal mCherry-tagged MBL-1 have short lifespan compared to N2. See S1-S2 Tables for lifespan statistics. (D) The most significant terms from WormExp output for upregulated genes (135) in day 2 adult *mbl-1(tm1563)* mutants compared to N2. (E) The most significant terms from WormExp output for downregulated genes (235) in day 2 adult *mbl-1(tm1563)* mutants compared to N2. In (D) and (E) numbers next to bars represent number of genes associated with the particular WormExp term. See S4 Table and S5 Table for differentially expressed genes between N2 and *mbl-1(tm1563)* mutants and WormExp statistics, respectively.

Since the reduced activity of MBNLs is a hallmark of DM1, we focused on the mechanism by which the loss of MBL-1 shortens lifespan. As an initial step, we performed RNA-seq of day 2 adult (day 5 from hatch) N2 and *mbl-1(tm1563)* mutants. Our analysis revealed that 135 genes are up- and 235 downregulated (at least 2-fold change in expression, FDR < 0.05) in *mbl-1(tm1563)* mutants compared to N2 (S4 Table). In order to identify aging-associated signaling pathways linking the loss of MBL-1 with shortened lifespan, we utilized the WormExp database, which collates nearly all published *C. elegans* expression data sets from public databases (YANG *et al*. 2016). We found that genes differentially regulated in *mbl-1(tm1563)* mutants are modulated by factors such as PQM-1, ELT-2, DAF-16 and PMK-1 (Fig 1D-1E and S5 Table). Interestingly, PMK-1, which encodes the conserved p38 mitogen-activated protein kinase (p38 MAPK), was the only factor associated exclusively with downregulated genes (Fig 1E and S5 Table). PMK-1-regulated genes, whose expression is reduced in day 15 vs. day 6 old *C. elegans* (YOUNGMAN *et al*. 2011), are also enriched among the downregulated genes in *mbl-1(tm1563)* mutants (Fig 1E and S5 Table). Due to this enrichment, we focused on the link between p38 MAPK/PMK-1 and MBL-1 to regulate lifespan.

### Loss of MBL-1 does not impair innate immune response

When subjecting up- and downregulated genes to gene ontology (GO) analysis by using the DAVID database (HUANG DA *et al*. 2009), we found that innate immune response-related GO terms are enriched among genes upregulated in *mbl-1(tm1563)* mutants (Fig 2A). This is interesting since the WormExp analysis indicated that PMK-1, a central factor of innate immunity-regulating p38 MAPK signaling (KIM *et al*. 2002), has reduced activity in *mbl-1(tm1563)* mutants (Fig 1E, S5 Table). On the other hand, WormBase GO enrichment analysis of 34 PMK-1-regulated genes whose expression is decreased in *mbl-1(tm1563)* mutants (Fig 1E, S5 Table) reveals that only 6 genes are associated with innate immunity (S5 Table). These data indicate that the loss of MBL-1 does not impair the expression of PMK-1-regulated innate immunity genes. Nevertheless, we asked whether the loss of MBL-1 affects pathogen resistance, and for this experiment we investigated the survival of *mbl-1(tm1563)* mutants on pathogenic *Pseudomonas aeruginosa* (PA14) in a slow-killing assay. In this assay we cultured PA14 on normal NGM plates where the killing occurs over the course of several days (TAN *et al*. 1999). *mbl-1(tm1563)* mutants were found to have similar survival on PA14 when compared to N2. This is in contrast to *pmk-1(km25)* mutants, which are extremely susceptible to PA14 (Fig 2B and S1-S2 Tables). Furthermore, *mbl-1(tm1563)* mutants show significantly elevated levels of phosphorylated p38 MAPK/PMK-1 upon PA14 infection (Fig 2C-2D, Fig S3 and S6 Table). The data demonstrate that MBL-1 dysfunction does not impair the ability to mount a p38 MAPK/PMK-1-mediated innate immune response upon infection.

**Fig 2.**
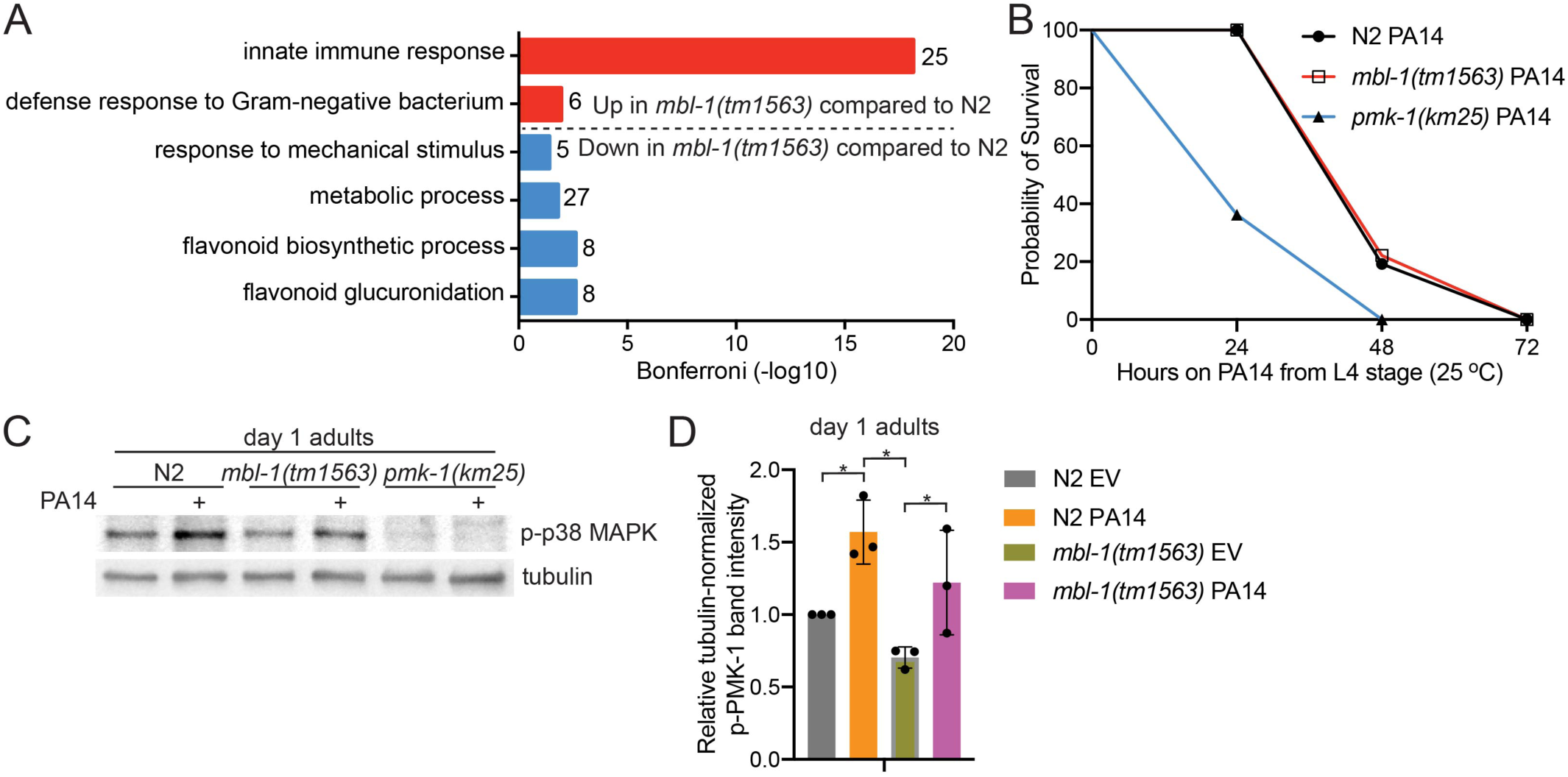
*mbl-1(tm1563)* mutants have innate immunity-associated gene expression signature, but normal response to *Pseudomonas aeruginosa*. (A) Gene Ontology (GO) Terms (biological process) enriched among genes that are up- and downregulated in day 2 adult *mbl-1(tm1563)* mutants compared to N2 in RNA-seq (GSE146801). Numbers next to bars represent number of genes associated with the particular GO term. See S4 Table for differentially expressed genes between N2 and *mbl-1(tm1563)* mutants. (B) *mbl-1(tm1563)* mutants have similar survival on pathogenic *Pseudomonas aeruginosa* (PA14) compared to N2, but enhanced survival compared to *pmk-1(km25)* mutants. See S1-S2 Tables for lifespan statistics. (C) Both N2 and *mbl-1(tm1563)* mutants have significantly elevated level of phosphorylated p38 MAPK upon PA14 infection. Animals were transferred to PA14 at L4 stage and collected 24 hours later (day 1 adults). (D) Quantified levels of phosphorylated p38 MAPK upon exposure to PA14. Bars represent the level of tubulin-normalized phosphorylated p38 MAPK relative to N2 EV with error bars indicating mean ± s.d. of three biological replicates (*p < 0.05, two-way ANOVA with Tukey’s test). See Fig S3 for Western blot repeats and S6 Table for Western blot quantifications.

### *mbl-1(tm1563)* mutants have reduced level of activated p38 MAPK/PMK-1, which contributes to their short lifespan

Since our RNA-seq data suggested that *mbl-1(tm1563)* mutants exhibit decreased PMK-1 activity (Fig 1E, S5 Table), we analyzed whether the level of phosphorylated p38 MAPK/PMK-1, which is traditionally used to assess the activity of this kinase, is altered in these mutants. Western blot analysis showed that day 2 old *mbl-1(tm1563)* mutants have reduced level of phosphorylated PMK-1 (Fig 3A-3B, Fig S3-S4 and S6 Table), thus demonstrating that basal p38 MAPK/PMK-1 activity is decreased upon disrupted MBL-1 function. *mbl-1* RNAi does not significantly affect PMK-1 phosphorylation (Fig 3C-3D, Fig S3 and S6 Table), which could be due to the insufficient knockdown efficiency. However, the quantified mean of phosphorylated PMK-1 level from three independent experiments (Fig 3C) indicates that also *mbl-1* knockdown modulates PMK-1 phosphorylation. Since MBNL1 is known to also regulate mRNA stability (MASUDA *et al*. 2012), we asked whether mRNA levels of *C. elegans* p38 MAPK pathway genes are altered in *mbl-1(tm1563)* mutants. We found that at the L4 larval stage the expression of *tir-1* is upregulated in *mbl-1(tm1563)* mutants, whereas the expression of *nsy-1*, *sek-1* and *pmk-1* is unchanged (Fig 3E, S7 Table). In day 2 adult animals only the expression of *nsy-1* is decreased (Fig 3F, S7 Table). As a conclusion, expression changes in p38 MAPK pathway genes are unlikely to explain the reduced p38 MAPK/PMK-1 activity in *mbl-1(tm1563)* mutants.

**Fig 3.**
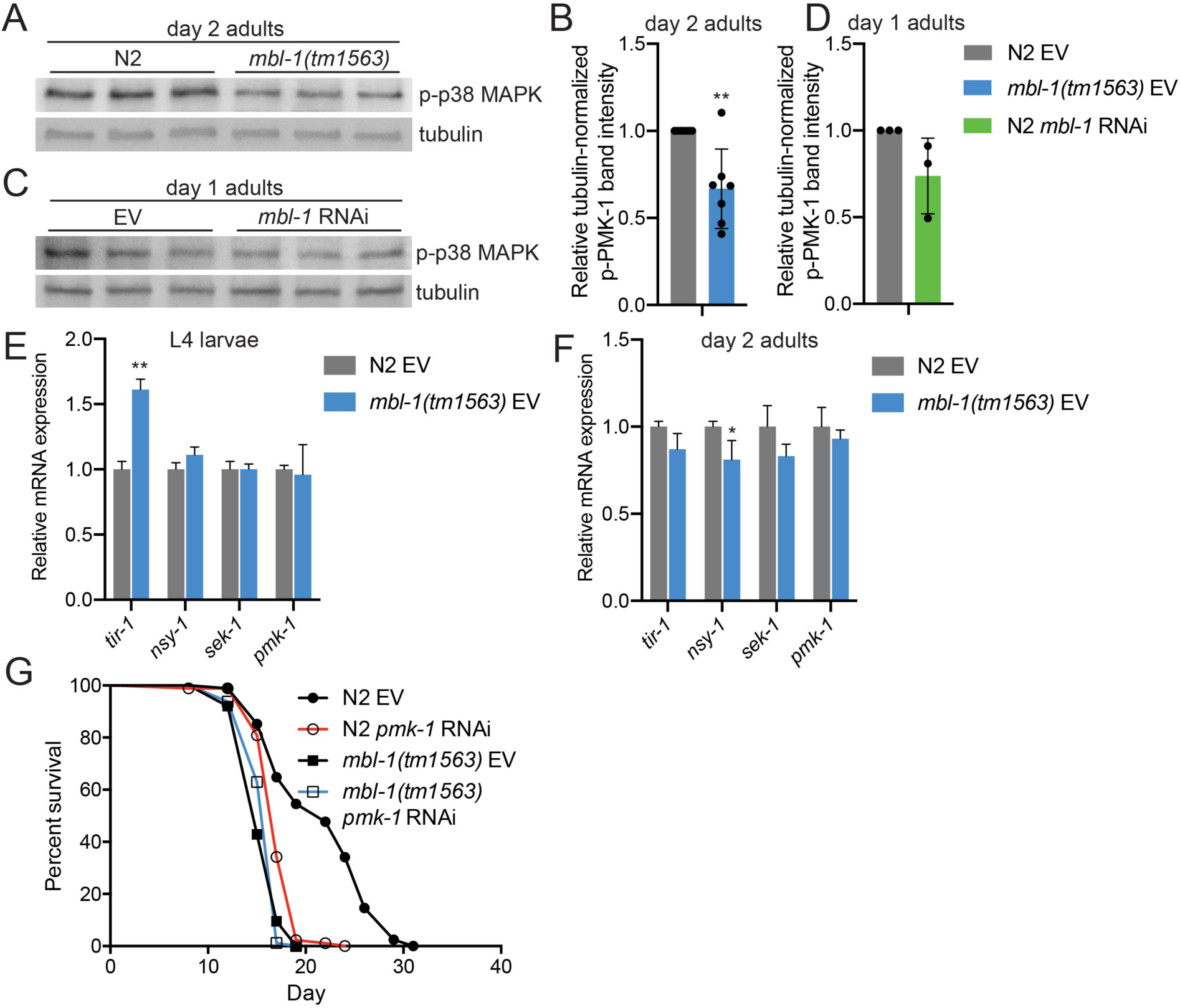
Decreased p38 MAPK/PMK-1 activation at non-pathogenic conditions shortens *mbl-1(tm1563)* mutant lifespan. (A) Day 2 adult *mbl-1(tm1563)* mutants have reduced level of phosphorylated p38 MAPK compared to N2. Shown are technical replicates of one biological replicate. (B) Quantified level of phosphorylated p38 MAPK in day 2 adult N2 and *mbl-1(tm1563)* mutants. (C) *mbl-1* RNAi does not significantly reduce the level of phosphorylated p38 MAPK in day 1 adult N2. (D) Quantified levels of phosphorylated p38 MAPK upon *mbl-1* RNAi in day 1 adult N2. In (B) and (D) bars represent the level of tubulin-normalized phosphorylated p38 MAPK relative to EV-treated N2 with error bars indicating mean ± s.d. of (B) seven and (D) three biological replicates (**p < 0.01, unpaired Student’s *t*-test). See Fig S3-S4 for Western blot repeats and S6 Table for Western blot quantifications. (E) The expression of p38 MAPK pathway components in L4 larvae N2 and *mbl-1(tm1563)* mutants. (F) The expression of p38 MAPK pathway components in day 2 adult N2 and *mbl-1(tm1563)* mutants. In (E) and (F) bars represent mRNA levels relative to N2 EV with error bars indicating mean ± s.d. of three biological replicates, each with three technical replicates (*p < 0.05, **p < 0.01, unpaired Student’s *t*-test). See S7 Table for raw qRT-PCR data. (G) *pmk-1* RNAi causes reduction in N2 lifespan (p < 0.01) but does not affect the lifespan of *mbl-1(tm1563)* mutants (p = 0.605). See S1-S2 Tables for lifespan statistics.

To determine whether the decreased p38 MAPK/PMK-1 activity contributes to the short lifespan of *mbl-1(tm1563)* mutants, we studied how these mutants are affected by the knockdown of *pmk-1.* RNAi of *pmk-1* reduces N2 lifespan, but does not affect the lifespan of *mbl-1(tm1563)* mutants (Fig 3G and S1-S2 Tables). Next we tested how RNAi of other components of the p38 MAPK pathway (Fig S5A) affect *mbl-1(tm1563)* mutant lifespan. We found that RNAi of *tir-1* shortens N2 lifespan but does not affect the lifespan of *mbl-1(tm1563)* mutants (Fig S5B, S1-S2 Tables). However, RNAi of *nsy-1* and *sek-1* does not affect N2 lifespan, whereas *sek-1* RNAi shortens *mbl-1(tm1563)* mutant lifespan (Fig S5C, S1-S2 Tables). These data indicate that the dysfunction of p38 MAPK/PMK-1 contributes to the short lifespan of *mbl-1(tm1563)* mutants independently of the upstream p38 MAPK signaling pathway.

### Reduced activity of ATF-7 and SKN-1 transcription factors modulate *mbl-1(tm1563)* mutant lifespan

In mammals, p38 MAPK signaling modulates cellular processes such as inflammation, development, cell differentiation and senescence by controlling the activity of multiple transcriptional regulators (ZARUBIN AND HAN 2005). Therefore, it is likely that reduced activity of p38 MAPK/PMK-1 kinase shortens the lifespan of *mbl-1(tm1563)* mutants by affecting the function of transcription factors. To this end, we investigated how downregulation of known PMK-1-regulated transcriptional regulators affect the lifespan of *mbl-1(tm1563)* mutants. Knockdown of *atf-7*, which encodes a transcription factor phosphorylated by PMK-1 in innate immune response (SHIVERS *et al*. 2010), induces larger decrease in N2 lifespan (14 %) than in *mbl-1(tm1563)* mutant lifespan (5 %) (Fig 4A and S1-S2 Tables). This finding is further supported by Cox proportional hazard regression analysis, which shows that *atf-7* RNAi increases hazard ratio (HR) more in N2 background than in *mbl-1(tm1563)* background (HR compared to N2 EV: N2 *atf-7* RNAi 2.290, *mbl-1(tm1563)* EV 3.728, *mbl-1(tm1563) atf-7* RNAi 4.742; HR compared to *mbl-1(tm1563)* EV: *mbl-1(tm1563) atf-7* RNAi 1.272). Since PMK-1 has been shown to phosphorylate and activate also SKN-1 (INOUE *et al*. 2005), we asked whether the aberrant function of this mammalian Nrf ortholog is linked with the shortened lifespan of *mbl-1(tm1563)* mutants. Similarly to *atf-7* knockdown, *skn-1* RNAi causes 1.8 % decrease in *mbl-1(tm1563)* mutant lifespan, whereas it shortens N2 lifespan by 14 % (HR compared to N2 EV: N2 *skn-1* RNAi 3.465, *mbl-1(tm1563)* EV 4.857, *mbl-1(tm1563) skn-1* RNAi 6.397; HR compared to *mbl-1(tm1563)* EV: *mbl-1(tm1563) skn-1* RNAi 1.317) (Fig 4B and S1-S2 Tables). To study further whether the loss of MBL-1 modulates SKN-1 function, we crossed *mbl-1(tm1563)* mutants with the transcriptional reporter of *gst-4* (*gst-4p::gfp*), which is a well-known target of SKN-1 (TULLET *et al*. 2008). The *mbl-1(tm1563)* mutants were found to have impaired induction of *gst-4p::gfp* expression upon oxidative stress (5 % H2O2 for 20 minutes, imaging after five hours recovery) (Fig 4C-4D, S7 Table), thus further demonstrating that SKN-1 activity is decreased in *mbl-1(tm1563)* mutants. Despite the requirement of MBL-1 on *gst-4* expression upon H2O2 treatment, MBL-1 expression pattern is not responsive to oxidative stress (Fig S1B).

**Fig 4.**
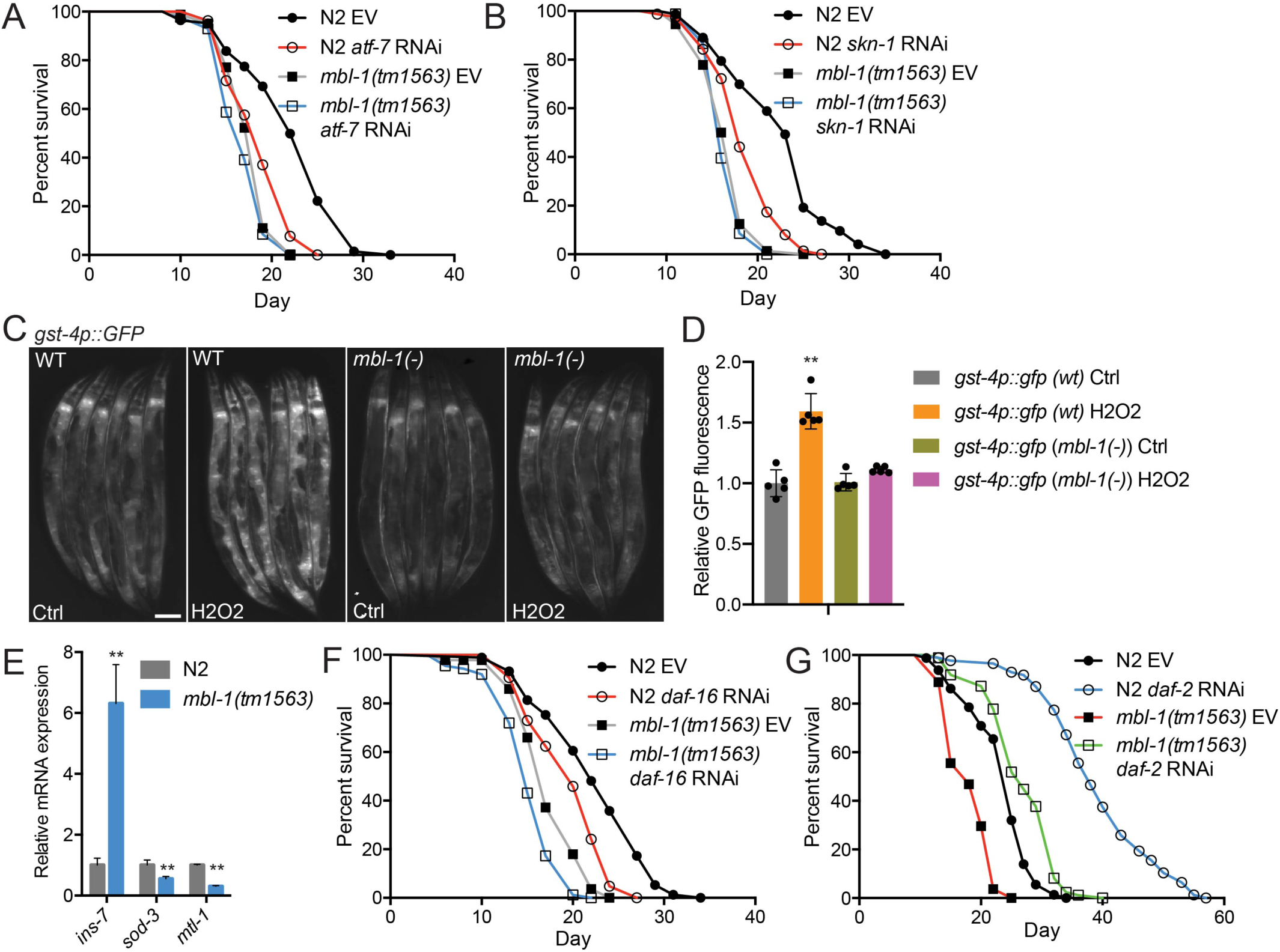
The short lifespan of *mbl-1(tm1563)* mutants is due to the dysfunction of ATF-7 and SKN-1 transcription factors. (A) *atf-7* RNAi shortens both N2 and *mbl-1(tm1563)* mutant lifespan (p < 0.01, p = 0.0291, respectively). (B) *skn-1* RNAi shortens both N2 and *mbl-1(tm1563)* mutant lifespan (p < 0.01, p = 0.0151, respectively). (C) *mbl-1(tm1563)* mutants crossed with the transcriptional *gst-4p::GFP* reporter strain (CL2166) show reduced fluorescence upon acute oxidative stress. Scale bar, 100 μm. (D) Quantification of *gst-4p::GFP* reporter strain fluorescence. Quantification was done in groups of six animals, n = 30 animals for each condition. Bars represent GFP fluorescence relative to wild-type (wt) Ctrl with error bars indicating mean ± s.d. of five replicates (*p < 0.05, one-way ANOVA with Tukey’s test). (E) Expression of DAF-16 regulated genes in N2 and *mbl-1(tm1563)* mutants. Animals were grown on EV and collected for qRT-PCR analysis at day 2 adult stage. Bars represent mRNA levels relative to N2 with error bars indicating mean ± s.d. of three biological replicates, each with three technical replicates (**p < 0.01, unpaired Student’s *t*-test). See S7 Table for raw qRT-PCR data. (F) *daf-16* RNAi shortens both N2 (p < 0.01) and *mbl-1(tm1563)* mutant (p < 0.01) lifespan. (G) *daf-2* RNAi increases both N2 (p < 0.01) and *mbl-1(tm1563)* mutant (p < 0.01) lifespans. See S1-S2 Tables for lifespan statistics.

The expression of *gst-4* is regulated also by DAF-16 (TULLET *et al*. 2008), the sole FOXO ortholog in *C. elegans*. DAF-16 is regulated by insulin/IGF-1-like signaling, which is a well-known pathway affecting aging (KENYON 2010). In addition to *gst-4*, our RNA-seq revealed that the expression of a sub-set of DAF-16/FOXO target genes is altered in *mbl-1(tm1563)* mutants (Fig 1D-1E and S4 Table). To further examine this, we analyzed mRNA levels of a few DAF-16 target genes by qRT-PCR in *mbl-1(tm1563)* mutants. We found that the expression of *sod-3* and *mtl-1*, which is promoted by DAF-16 (MURPHY *et al*. 2003), is downregulated in *mbl-1(tm1563)* mutants, whereas the expression of *ins-7*, which is suppressed by DAF-16 (MURPHY *et al*. 2003), is upregulated upon the loss of MBL-1 (Fig 4E). Despite these changes in DAF-16 target gene expression, longevity assays showed that *daf-16* RNAi shortens N2 and *mbl-1(tm1563)* mutant lifespan by 6.8 % and 9 %, respectively (Fig 4F and S1-S2 Tables), consistent with the notion that the loss of MBL-1 shortens lifespan independently of DAF-16. Additionally, RNAi of *daf-2*, which leads to increased longevity in an DAF-16-dependent manner (KENYON *et al*. 1993), results in a similar increase in both N2 and *mbl-1(tm1563)* mutant lifespan (31.9 % and 32.7 %, respectively) (Fig 4G and S1-S2 Tables). Taken together, these data indicate that loss of MBL-1 leads to shortened lifespan by reducing the activity of p38 MAPK/PMK-1, which in turn decreases the activity of ATF-7 and SKN-1 transcription factors. Furthermore, although the loss of MBL-1 causes differential expression of some DAF-16 target genes, the altered DAF-16 function does not contribute to the short lifespan of *mbl-1(tm1563)* mutants.

### Mitochondrial stress rescues the short lifespan of *mbl-1(tm1563)* mutants

Next, we asked whether targeting the known aging mechanisms could modulate p38 MAPK activity, and thereby, rescue the short lifespan of *mbl-1(tm1563)* mutants. Interestingly, it has been shown that inhibition of mitochondrial respiratory chain complex I activates the PMK-1/ATF-7 signaling pathway (CHIKKA *et al*. 2016). Moreover, mitochondrial chaperone HSP-60 binds and stabilizes SEK-1 MAPK (see Fig S5A for *C. elegans* p38 MAPK pathway), thus promoting PMK-1 activation (JEONG *et al*. 2017). Due to these reports, we asked whether mitochondrial function could extend *mbl-1(tm1563)* mutant lifespan.

To study a role for mitochondrial stress on the *mbl-1(tm1563)* mutant lifespan, we used RNAi of mitochondrial respiratory chain complex IV subunit *cox-5B* (also known as *cco-1*), which is known to extend lifespan (DILLIN *et al*. 2002). As *cox-5B* RNAi also promotes innate immunity (MATILAINEN *et al*. 2017), we hypothesized that it leads to activation of p38 MAPK/PMK-1 kinase. Strikingly, we found that *cox-5B* RNAi leads to a significantly larger (p < 0.01, Cox proportional hazard regression analysis) lifespan extension in *mbl-1(tm1563)* mutants (+31 %) compared to N2 animals (+15.3 %) (HR compared to N2 EV: N2 *cox-5B* RNAi 0.42, *mbl-1(tm1563)* EV 3.426, *mbl-1(tm1563) cox-5B* RNAi 0.302; HR compared to *mbl-1(tm1563)* EV: *mbl-1(tm1563) cox-5B* RNAi 0.088) (Fig 5A and S1-S2 Tables). To test whether *cox-5B* RNAi affects the p38 MAPK pathway, we performed Western blot to analyze p38 MAPK/PMK-1 phosphorylation. We found that *cox-5B* knockdown does not increase the level of phosphorylated p38 MAPK/PMK-1 in L3 larvae (Fig 5B-5C, Fig S4, S6 Table). At L4 larval stage *cox-5B* RNAi increases the level of phosphorylated p38 MAPK/PMK-1 in both N2 and *mbl-1(tm1563)* mutants in all individual experiments (Fig 5B, Fig S4, S6 Table), but due to the variation between experiments, the increase is not statistically significant (Fig 5D). Also in day 2 adults the effect of *cox-5B* RNAi on p38 MAPK/PMK-1 phosphorylation varies between experiments, and the increase in p38 MAPK/PMK-1 phosphorylation is not statistically significant (Fig 5E-5F, Fig S4, S6 Table). However, in day 2 adult *mbl-1(tm1563)* mutants *cox-5B* RNAi increases the level of phosphorylated p38 MAPK/PMK-1 in four out of five independent experiments (Fig S4, S6 Table), thus indicating that similar to L4 larvae, *cox-5B* RNAi may modulate the activation p38 MAPK/PMK-1 in day 2 adults.

**Fig 5.**
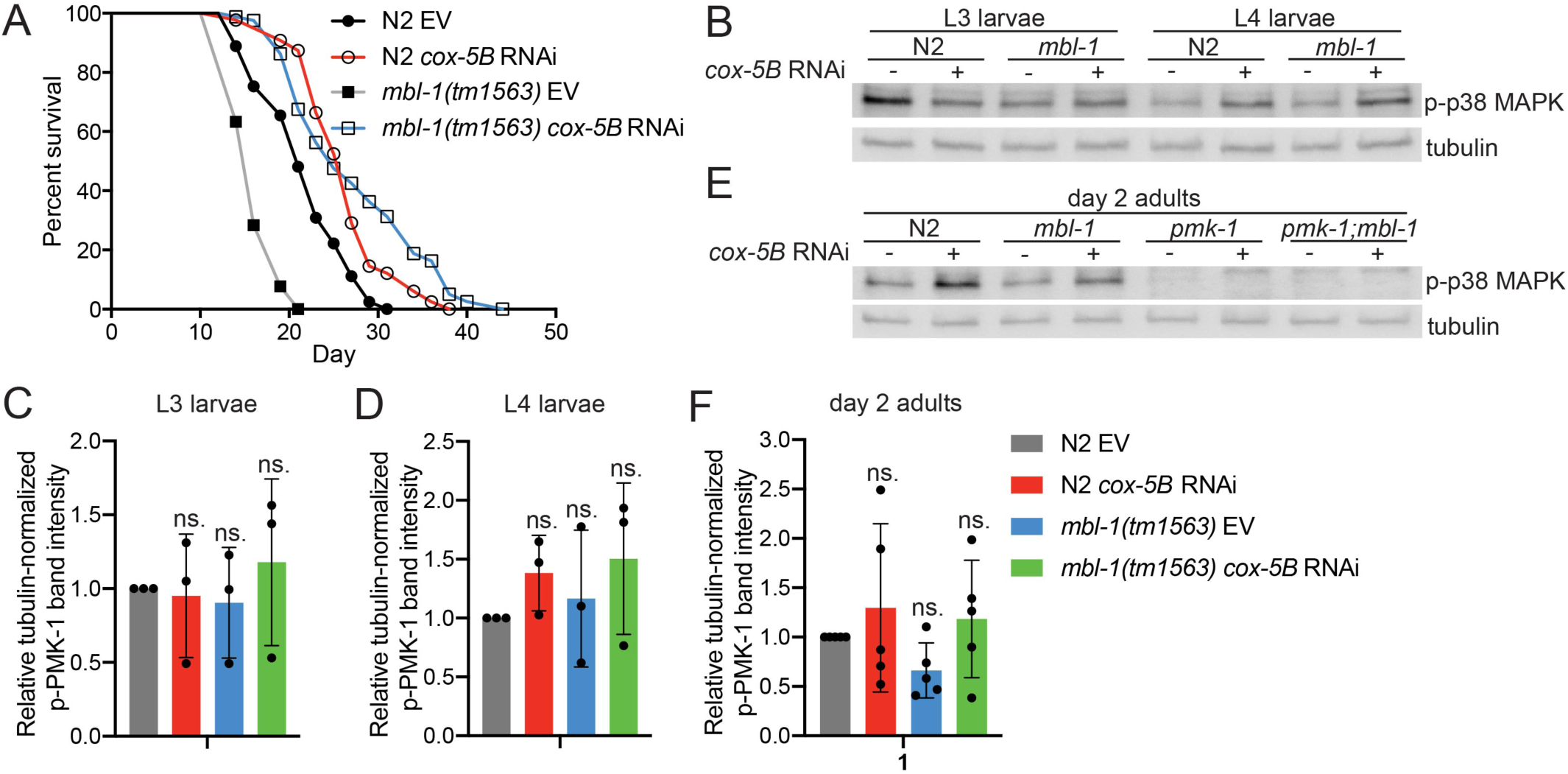
*cox-5B* RNAi-mediated mitochondrial stress modulates the level of phosphorylated p38 MAPK/PMK-1 and leads to increase in *mbl-1(tm1563)* mutant lifespan. (A) *cox-5B* RNAi-treated *mbl-1(tm1563)* mutants have longer lifespan than *cox-5B* RNAi-treated N2 animals (p < 0.01). (B) Western blot of phosphorylated p38 MAPK/PMK-1 in L3 and L4 larvae N2 and *mbl-1(tm1563)* mutants upon *cox-5B* RNAi. (C-D) Quantified levels of phosphorylated p38 MAPK upon *cox-5B* RNAi in (C) L3 and (D) L4 larvae N2 and *mbl-1(tm1563)* mutants. (E) Western blot of phosphorylated p38 MAPK/PMK-1 in day 2 adult N2 and *mbl-1(tm1563)* mutants upon *cox-5B* RNAi. (F) Quantified levels of phosphorylated p38 MAPK upon *cox-5B* RNAi in day 2 adult N2 and *mbl-1(tm1563)* mutants. In (C), (D) and (F) bars represent the level of tubulin-normalized phosphorylated p38 MAPK relative to EV-treated N2 with error bars indicating mean ± s.d. of (C-D) three and (F) five biological replicates (two-way ANOVA with Tukey’s test). See Fig S4 for Western blot repeats and S6 Table for Western blot quantifications.

Next we asked how *cox-5B* RNAi-mediated mitochondrial stress affects the expression of PMK-1-regulated genes (TROEMEL *et al*. 2006; YOUNGMAN *et al*. 2011). Importantly, the majority of PMK-1 target genes have decreased expression in *mbl-1(tm1563)* mutants (Fig 6A). The downregulated genes include C17H12.8 and K08D8.5 (Fig 6A), which are also regulated by ATF-7 (SHIVERS *et al*. 2010), providing further evidence for ATF-7 dysfunction in *mbl-1(tm1563)* mutants. Although less efficiently than *mbl-1(tm1563)* mutation, also *mbl-1* RNAi leads to decreased expression of a sub-set of PMK-1 target genes (Fig 6B). We found that *cox-5B* knockdown increases the expression of *mul-1*, *catp-3* and *amt-1* in both N2 and *mbl-1(tm1563)* mutants (Fig 6C). To examine whether the upregulation of these genes is dependent on PMK-1, we compared their expression between *mbl-1(tm1563)* mutants and *pmk-1(km25)*;*mbl-1(tm1563)* double mutants. Loss of PMK-1 blunts the *cox-5B* RNAi-mediated induction of these genes (Fig 6D). The data further demonstrate that *cox-5B* knockdown modulates p38 MAPK/PMK-1 activity.

**Fig 6.**
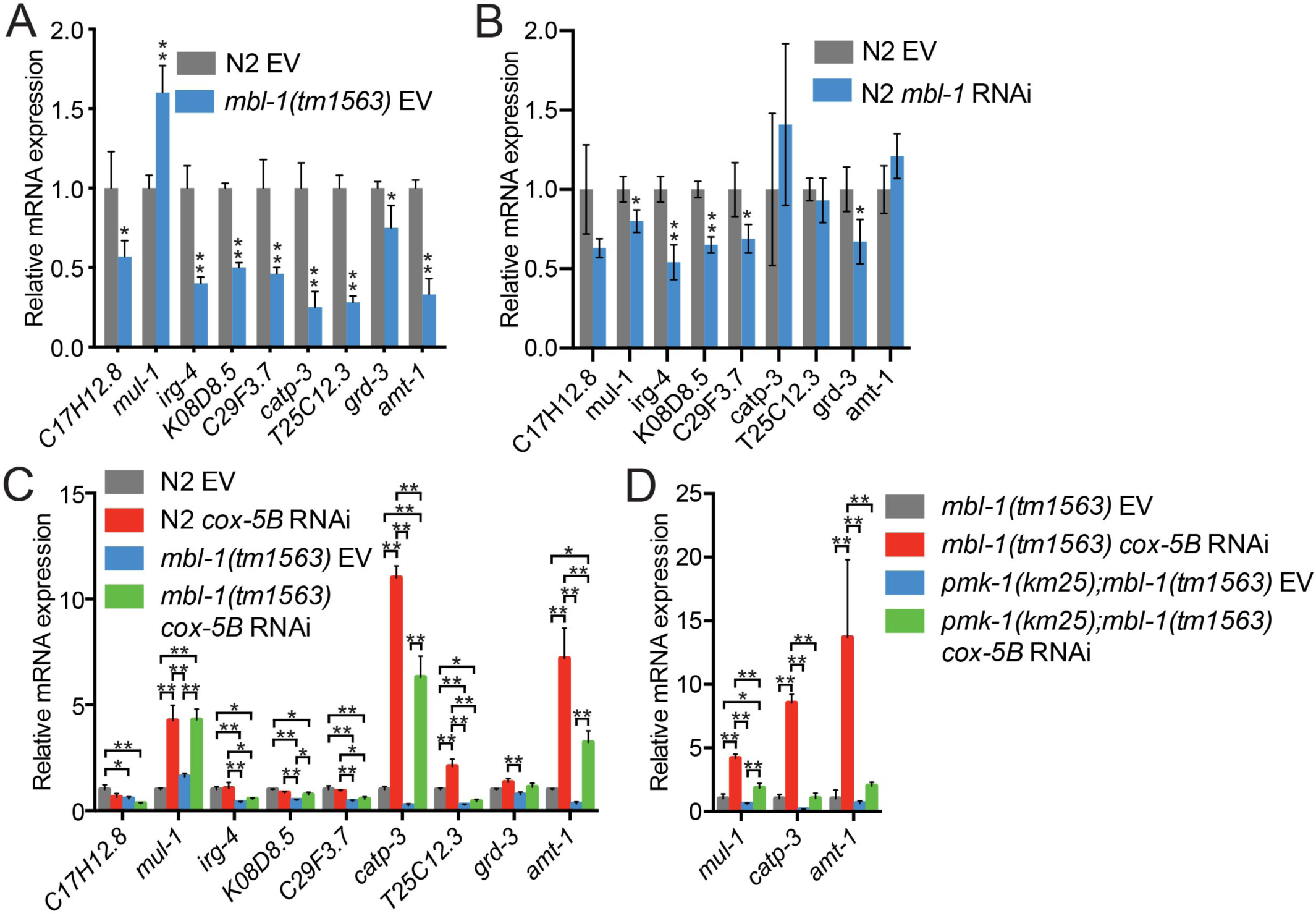
Loss of MBL-1 leads to altered *cox-5B* RNAi-mediated mitochondrial stress modulates the expression of p38 MAPK/PMK-1 target genes. **(A) The expression of PMK-1-regulated genes in L4 larvae N2 and *mbl-1(tm1563)* mutants. (B) The expression of PMK-1-regulated genes in *mbl-1* RNAi-treated L4 larvae N2. In (A) and (B) bars represent mRNA levels relative to N2 EV with error bars indicating mean ± s.d. of three biological replicates, each with three technical replicates (*p < 0.05, **p < 0.01, unpaired Student’s *t*-test). (C) The expression of PMK-1 regulated genes in L4 larvae N2 and *mbl-1(tm1563)* mutants upon *cox-5B* RNAi. The EV bars are from the same experiment presented in (A). (D) The expression of selected PMK-1 targets in L4 larvae *mbl-1(tm1563)* mutants and *pmk-1(km25);mbl-1(tm1563)* double mutants. In (C) and (D) bars represent mRNA levels relative to (C) N2 EV or (D) *mbl-1(tm1563)* EV with error bars indicating mean ± s.d. of three biological replicates, each with three technical replicates (*p < 0.05, **p < 0.01, two-way ANOVA with Tukey’s test). See S7 Table for raw qRT-PCR data.**

### *mbl-1(tm1563)* mutants require p38 MAPK/PMK-1 for maximal lifespan extension upon mitochondrial stress

Although mitochondrial stress induces innate immune response (PELLEGRINO *et al*. 2014; CHIKKA *et al*. 2016; JEONG *et al*. 2017; MATILAINEN *et al*. 2017), p38 MAPK/PMK-1 has not been associated with mitochondria-mediated longevity. We asked whether PMK-1 is required for the increased *mbl-1(tm1563)* mutant lifespan upon *cox-5B* RNAi. Although *cox-5B* RNAi extends the lifespan of all tested strains (Fig 7A-7B and S1-S2 Tables), *pmk-1(km25)*;*mbl-1(tm1563)* double mutation blocks the maximal lifespan extension observed with *cox-5B* RNAi-treated *mbl-1(tm1563)* mutants (Fig 7A-7B and S1-S2 Tables). Importantly, *mbl-1(tm1563)* mutants and *pmk-1(km25)*;*mbl-1(tm1563)* double mutants have similar lifespan on control RNAi (EV) (Fig 7A-7B and S1-S2 Tables). In addition to *cox-5B* RNAi, we also investigated whether PMK-1 is required for the lifespan increase of *mbl-1(tm1563)* mutants upon RNAi of mitochondrial respiratory chain complex I subunit *nduf-6*, which has been shown to increase p38 MAPK/PMK-1 phosphorylation (CHIKKA *et al*. 2016). Although on *nduf-6* RNAi *mbl-1(tm1563)* mutant lifespan is not increased beyond *nduf-6* RNAi-treated N2 animals, *nduf-6* knockdown leads to 30.6 % increase in *mbl-1(tm1563)* mutant lifespan (HR compared to *mbl-1(tm1563)* EV: 0.089), whereas N2 lifespan is increased by 15.7 % (HR compared to N2 EV: 0.507) (Fig S5D-S5E and S1-S2 Tables). Furthermore, similarly as with *cox-5B* RNAi, *pmk-1(km25)*;*mbl-1(tm1563)* double mutation blocks the maximal lifespan extension observed with *nduf-6* RNAi-treated *mbl-1(tm1563)* mutants (Fig S5D-S5E and S1-S2 Tables). Altogether, these data demonstrate that mitochondrial stress modulates the activity of p38 MAPK/PMK-1, thereby resulting in extended lifespan.

**Fig 7.**
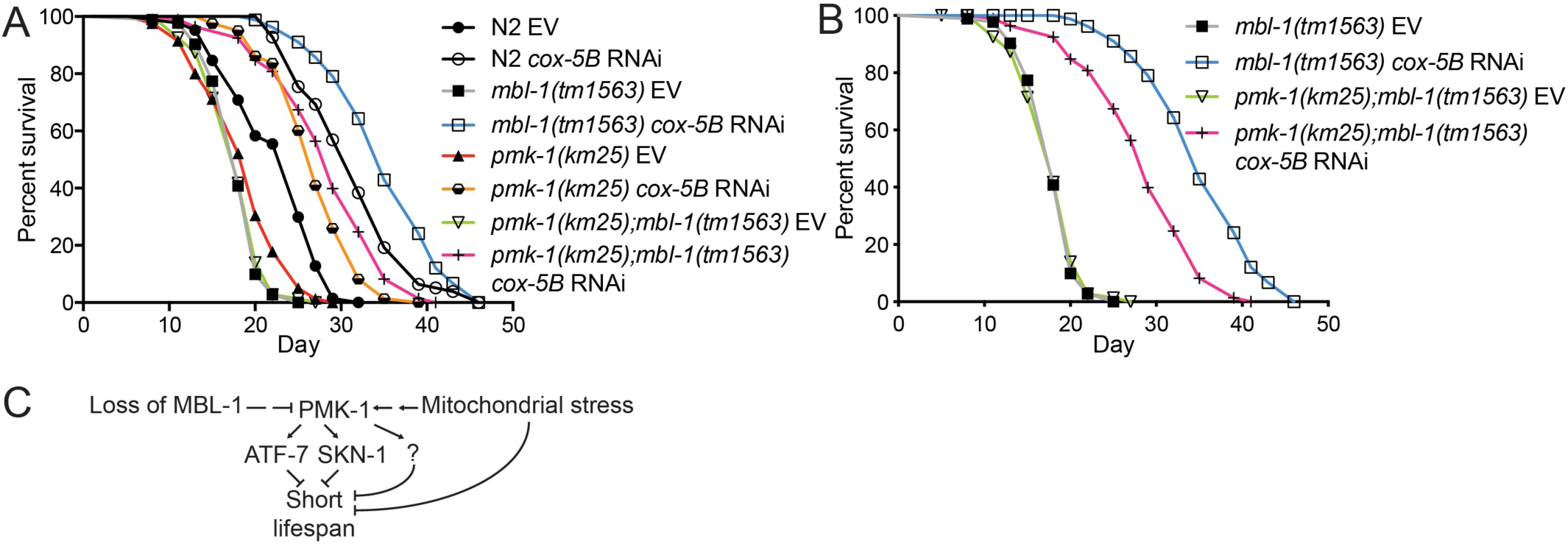
*cox-5B* RNAi-mediated mitochondrial stress results in PMK-1-dependent increase in *mbl-1(tm1563)* mutant lifespan. (A-B) *cox-5B* RNAi-treated *mbl-1(tm1563)* mutants have extended longevity compared to *cox-5B* RNAi-treated N2 (p < 0.01), *pmk-1(km25)* (p < 0.01) and *pmk-1(km25);mbl-1(tm1563)* (p < 0.01) mutants. (B) shows the sub-set of lifespan curves presented in (A). See S1-S2 Tables for lifespan statistics. (C) Model based on the data presented here.

## Discussion

The data presented here support the hypothesis that loss of MBL-1 disrupts the mechanism maintaining basal p38 MAPK/PMK-1 activity in normal conditions, which leads to reduced activity of transcription factors ATF-7 and SKN-1 and shortened lifespan (Fig 7C). Since the decline in basal p38 MAPK/PMK-1 activity is an aging-associated process (YOUNGMAN *et al*. 2011), it can be hypothesized that the aberrant p38 MAPK signaling contributes to the premature aging-phenotype observed with MBNL deficiency in DM1 patients. Although *mbl-1(tm1563)* mutants have a decreased level of activated p38 MAPK/PMK-1 at non-pathogenic condition, they do not show increased susceptibility to pathogens (Fig 2B and S1-S2 Tables). This demonstrates that p38 MAPK/PMK-1 regulates lifespan independently of its role in innate immunity. In this context, chronic stress caused by aging differs from acute stress caused by pathogen infection. Therefore, we propose that moderate reduction in p38 MAPK/PMK-1 activity shortens *mbl-1(tm1563)* mutant lifespan, but it retains sufficient activity to respond to acute pathogenic bacterial stress. On the other hand, upregulation of innate immunity-associated genes in *mbl-1(tm1563)* mutants (Fig 2A) suggests that other pathogen surveillance-mechanisms may compensate the reduced p38 MAPK/PMK-1 activity, thus ensuring the normal survival of *mbl-1(tm1563)* mutants on PA14 (Fig 2B).

Importantly, the role of p38 MAPK/PMK-1 as a regulator of lifespan under non-pathogenic conditions is controversial, as it has been shown that *pmk-1(km25)* mutants have either normal lifespan (KIM *et al*. 2002; TROEMEL *et al*. 2006; WU *et al*. 2019) or reduced lifespan (PUJOL *et al*. 2008; PARK *et al*. 2018; ZHOU *et al*. 2019). Since PMK-1 is required for response to biotic stimulus, one possible explanation for differences between laboratories could be due to bacterial growth conditions, which in turn may be affected by brand of peptone and agar used for plate media. Interestingly, although we found that both *pmk-1(km25)* mutants and *pmk-1* RNAi-treated animals have shortened lifespan, the hazard ratio is bigger for *pmk-1* RNAi-treated animals in Fig 3G (HR compared to N2 EV: 5.062) than for *pmk-1(km25)* mutants in Fig 7A (HR compared to N2 EV: 2.406) and in Fig S5D (HR compared to N2 EV: 2.769). Although conclusions should not be made by comparing independent lifespan experiments, these data suggest that acute loss of PMK-1 could be more detrimental relative to chronic PMK-1 depletion. To hypothesize further, it is possible that *pmk-1(km25)* mutants have activated stress responses, which in turn can lead to inconsistencies in lifespan experiments in different laboratory settings.

Although demonstrating that the MBL-1 alternative splicing factor is required to maintain the basal p38 MAPK/PMK-1 activity and normal lifespan, this study leaves an open question about the molecular mechanism behind this phenotype. Interestingly, in addition to alternative splicing, MBNL1 regulates also mRNA decay and mRNA localization (MASUDA *et al*. 2012; WANG *et al*. 2012). Hence, it is possible that MBL-1 affects p38 MAPK/PMK-1 activity through a combination of these mechanisms. Interestingly, p38 MAPK can also be activated by mechanical stimuli (NGUYEN *et al*. 2000; ZHANG *et al*. 2000; SAWADA *et al*. 2001; WANG *et al*. 2005; LEWTHWAITE *et al*. 2006; CHAUDHURI AND SMITH 2008; DOLINAY *et al*. 2008; HOFFMAN *et al*. 2017). Since *mbl-1* mutants display improper formation of neuromuscular junction synapses and altered locomotion (SPILKER *et al*. 2012), one intriguing hypothesis is that dysfunctional neuromuscular communication and the consequent abnormal muscle function leads to lower level of mechanical stimuli, and thus to decreased basal p38 MAPK/PMK-1 activity. In this context, one interesting experiment would be to address whether electrical stimulation promote basal p38 MAPK/PMK-1 activity and increase in lifespan in *mbl-1* mutants.

In addition to the model that MBL-1 promotes normal lifespan by maintaining the basal p38 MAPK/PMK-1 signaling (Fig 7C), our data show that mitochondrial stress promotes the longevity of *mbl-1(tm1563)* mutants in a partially PMK-1-dependent manner (Fig 7A-7B, Fig S5D-S5E and S1-S2 Tables). Although Western blot and qRT-PCR data (Fig 5B-5F, Fig S4, Fig 6C-D) indicate that *cox-5B* knockdown modulates the activity of p38 MAPK/PMK-1, it must be taken into account that p38 MAPK can also be activated by acetylation (PILLAI *et al*. 2011), raising the possibility that mitochondrial stress promotes p38 MAPK/PMK-1 function through different mechanisms. Importantly, overactive p38 MAPK signaling is toxic and shortens lifespan (CHEESMAN *et al*. 2016), and therefore, it can be assumed that p38 MAPK/PMK-1 is only moderately activated during non-infectious conditions such as mitochondrial stress. Since p38 MAPK/PMK-1 is only partially required for the increased *mbl-1(tm1563)* mutant lifespan upon mitochondrial stress (Fig 7A-7B, Fig S5D-S5E and S1-S2 Tables), there are also other mechanisms involved. For example, it is possible that the loss of MBL-1 disrupts mitochondrial function, and therefore, mitochondrial perturbation-activated stress responses may restore the homeostasis in this organelle, thus leading to strikingly increased lifespan.

Taken together, these data elucidate an important mechanism of how the loss of DM1-associated muscleblind splicing factor affects lifespan. In terms of quality of life, it has been proposed that targeting aging may be a more efficient approach compared to targeting individual diseases (KAEBERLEIN *et al*. 2015). Therefore, since DM1 is a multisystemic disease with similarities to aging (MATEOS-AIERDI *et al*. 2015; MEINKE *et al*. 2018), this study suggests that targeting the aging process could provide a powerful complementary therapeutic approach for this severe disorder.

## Acknowledgements

The Garcia laboratory is supported by the Academy of Finland Project Funding (309173) and by the Institute of Biotechnology, University of Helsinki. O.M. is supported by the University of Helsinki three-year grant. A.R.S.R. was supported by the INOV Contacto-program. R.I.S. is supported by NIH NIDDK P30DK040561. Some strains were provided by the CGC, which is funded by NIH Office of Research Infrastructure Programs (P40 OD010440). We also thank the National BioResource Project (NBRP) for sharing the *mbl-1(tm1563)* strain, the laboratory of Dr. Carina Holmberg for sharing reagents, and Garcia lab members and Dr. Brendan Battersby for comments on the manuscript.

## Supplemental material

**S1 Fig.**
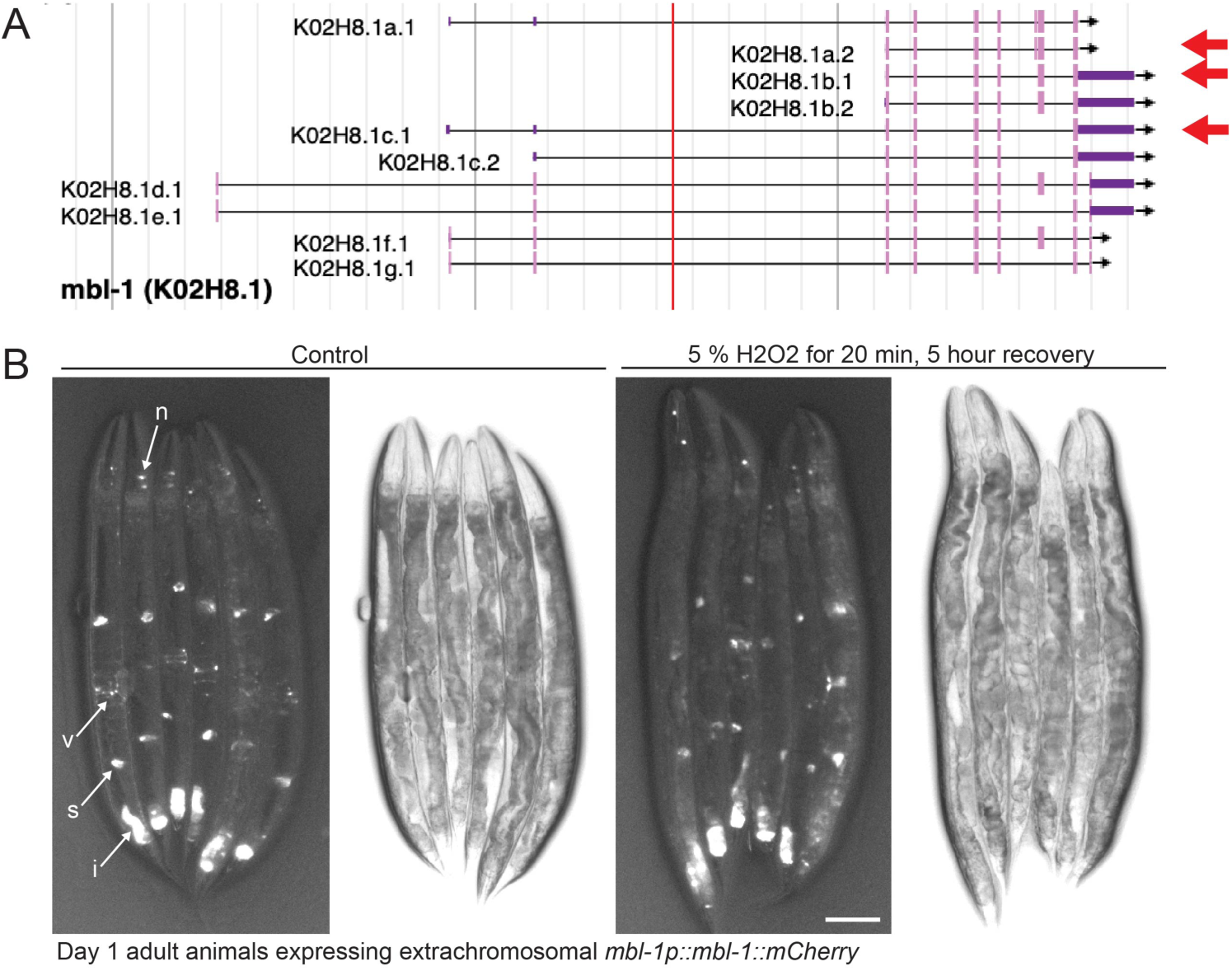
Overexpression of mCherry-tagged MBL-1. (A) Screenshot from WormBase presenting mbl-1 gene. Arrows indicate isoforms expressed in mbl-1 OE lines. (B) Day 1 adult animals expressing mbl-1p::mbl-1::mCherry under normal conditions (Ctrl, EV) and after acute oxidative stress (5 % H2O2 for 20 minutes, image taken after five hours recovery on EV). Abbreviations: n, neuron; v, vulva; s, spermatheca; i, intestine. Scale bar, 100 μm.

**S2 Fig.**
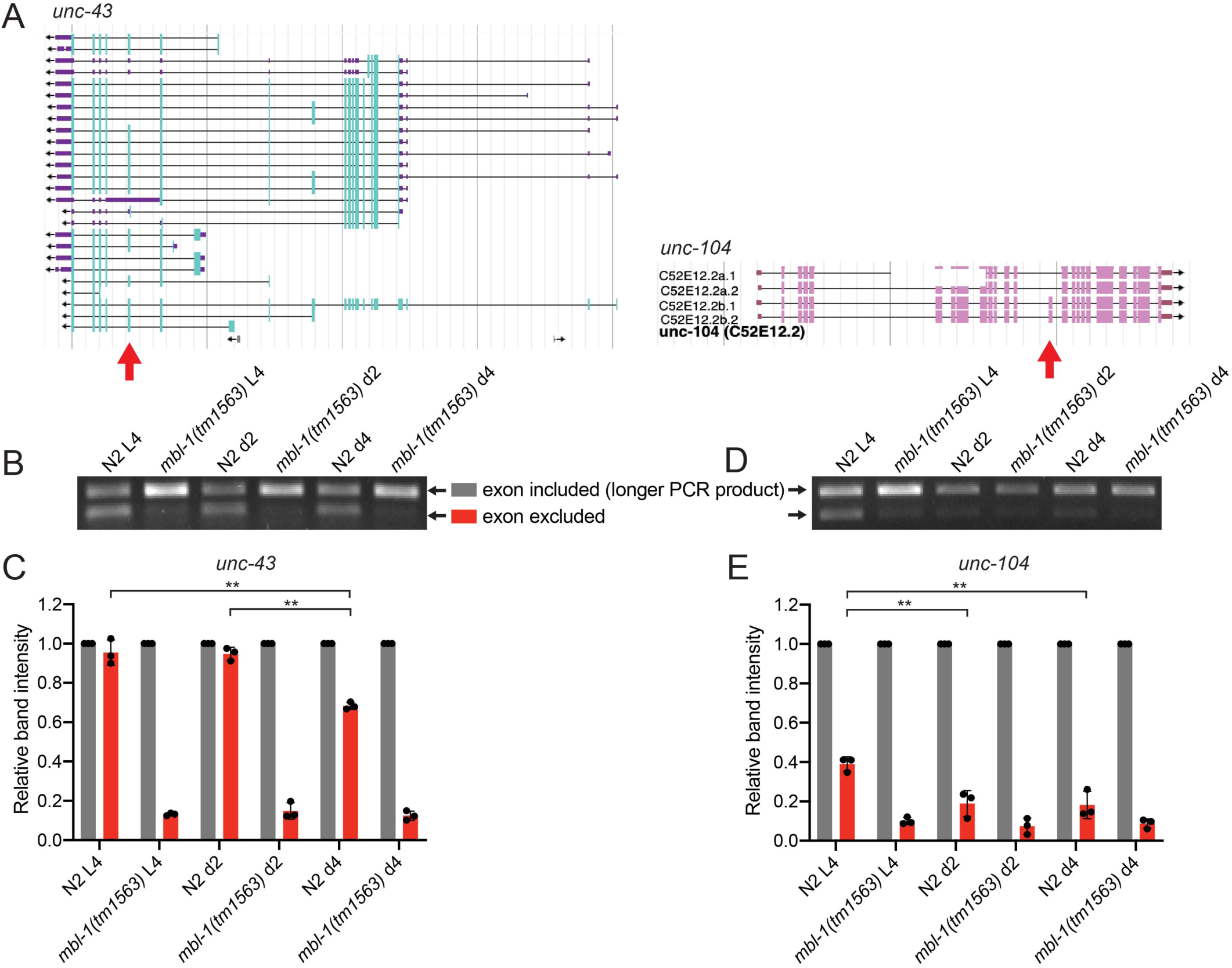
MBL-1 activity decreases upon aging. (A) Screenshot from WormBase presenting unc-43 and unc-104 genes. Arrows indicate exons whose exclusion were examined upon aging. (B-C) PCR-mediated splicing assay of unc-43 and (D-E) unc-104 genes. (B) and (D) show representative agarose gels and (C) and (E) agarose gel quantifications. In (C) and (E) bars represent the intensity of shorter PCR product (exon excluded) relative to longer PCR product (exon included) with error bars indicating mean ± s.d. of three biological replicates (**p < 0.01, two-way ANOVA with Tukey’s test). See S3 Table for agarose gel quantifications.

**S3 Fig.**
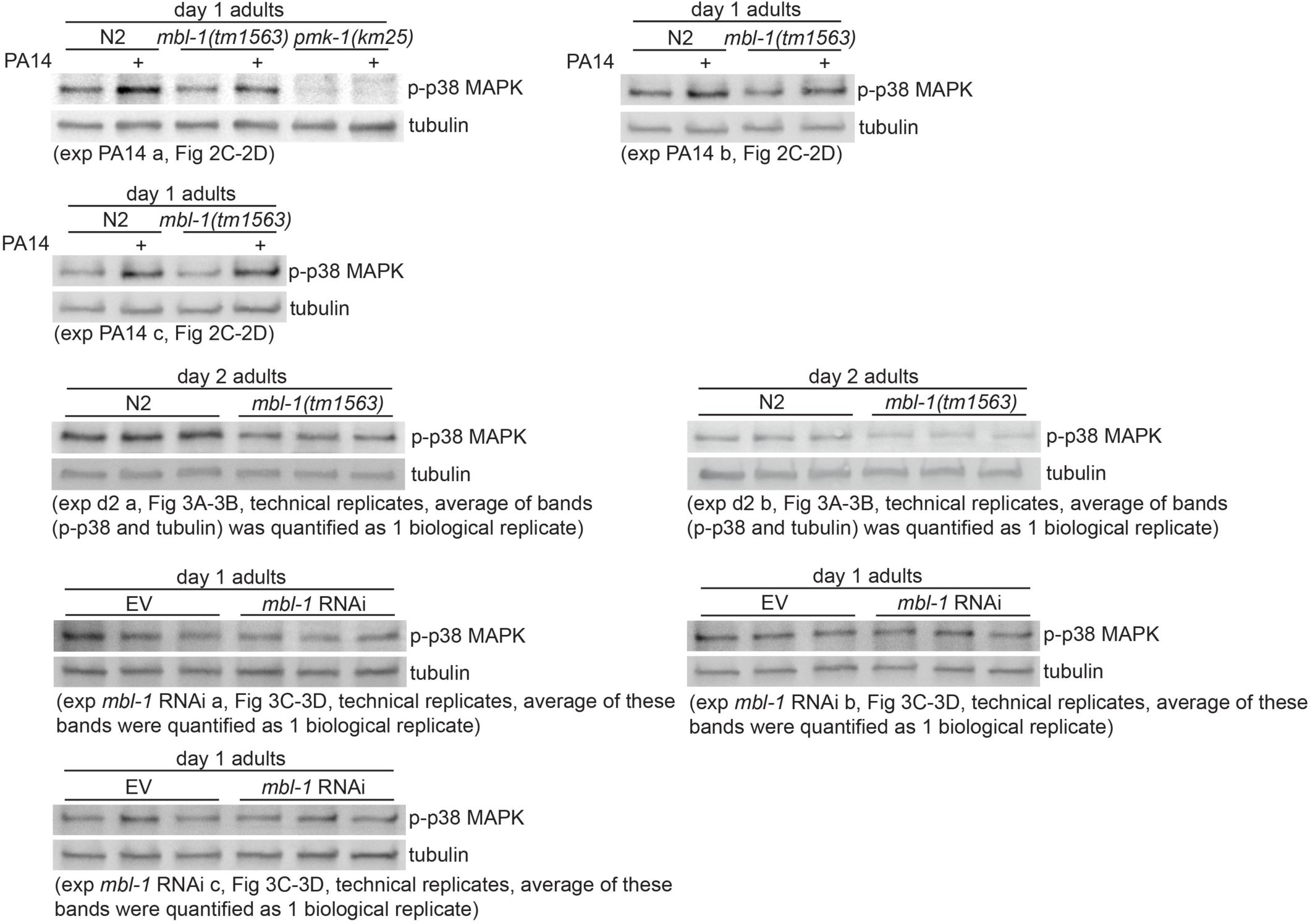
Western blot repeats for figures 2C, 3A and 3C. See S6 Table for Western blot quantifications.

**S4 Fig.**
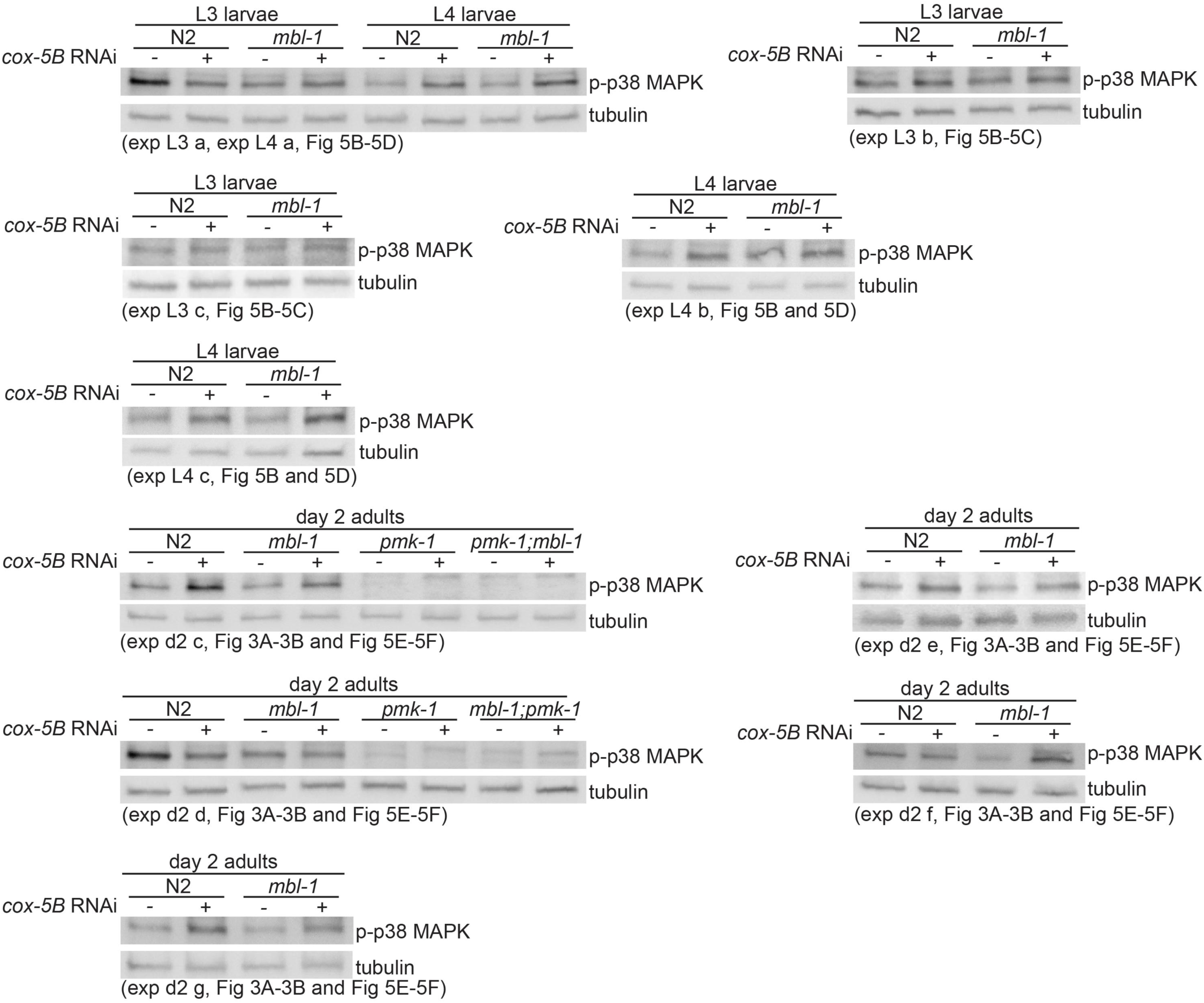
Western blot repeats for figures 3A, 5B and 5E. See S6 Table for Western blot quantifications.

**Fig S5.**
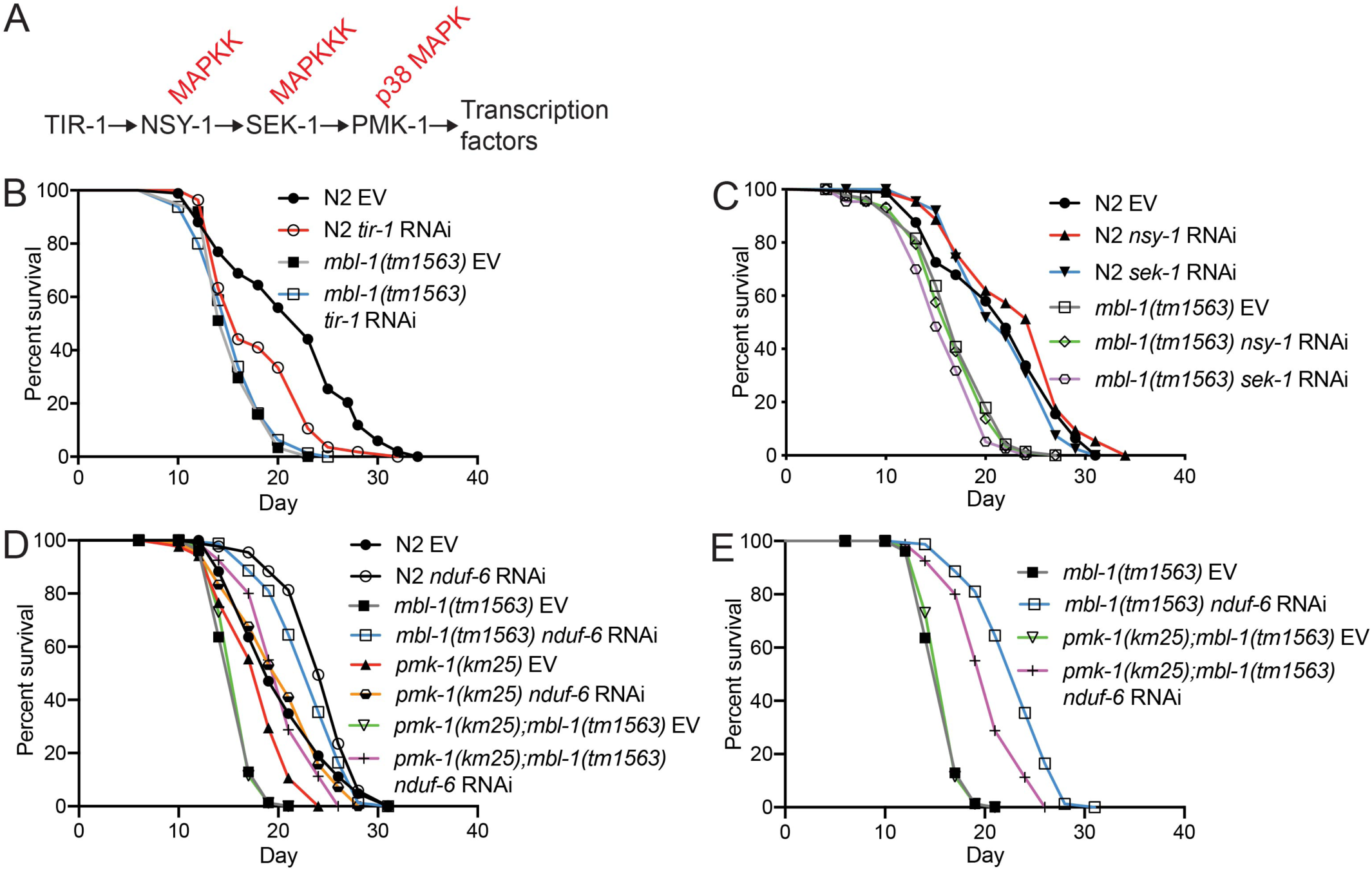
Lifespan effects upon knockdown of p38 MAPK pathway components *tir-1*, *nsy-1* and *sek-1* and mitochondrial electron transport chain subunit *nduf-6*. (A) *C. elegans* p38 MAPK pathway. (B) *tir-1* RNAi causes reduction in N2 lifespan, but does not affect the lifespan of *mbl-1(tm1563)* mutants. (C) *nsy-1* and *sek-1* RNAi does not affect the lifespan of N2, whereas *sek-1* RNAi shortens the lifespan of *mbl-1(tm1563)* mutants. (D-E) *nduf-6* RNAi-treated *mbl-1(tm1563)* mutants have extended longevity compared to *nduf-6* RNAi-treated *pmk-1(km25);mbl-1(tm1563)* mutants. (D) shows the sub-set of lifespan curves presented in (E). See S1-S2 Tables for lifespan statistics.

**S1 Table.**
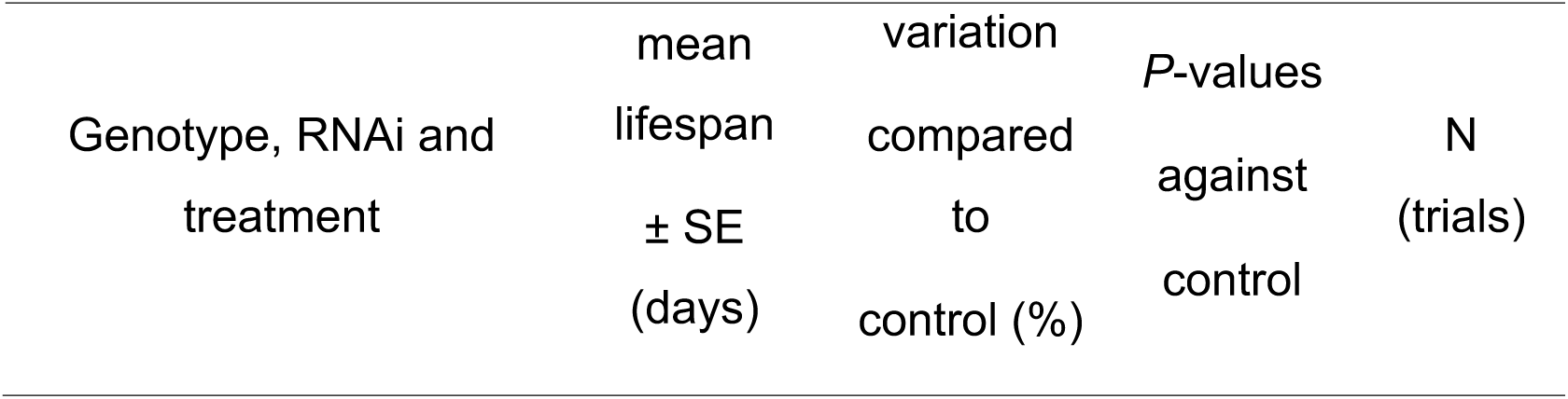
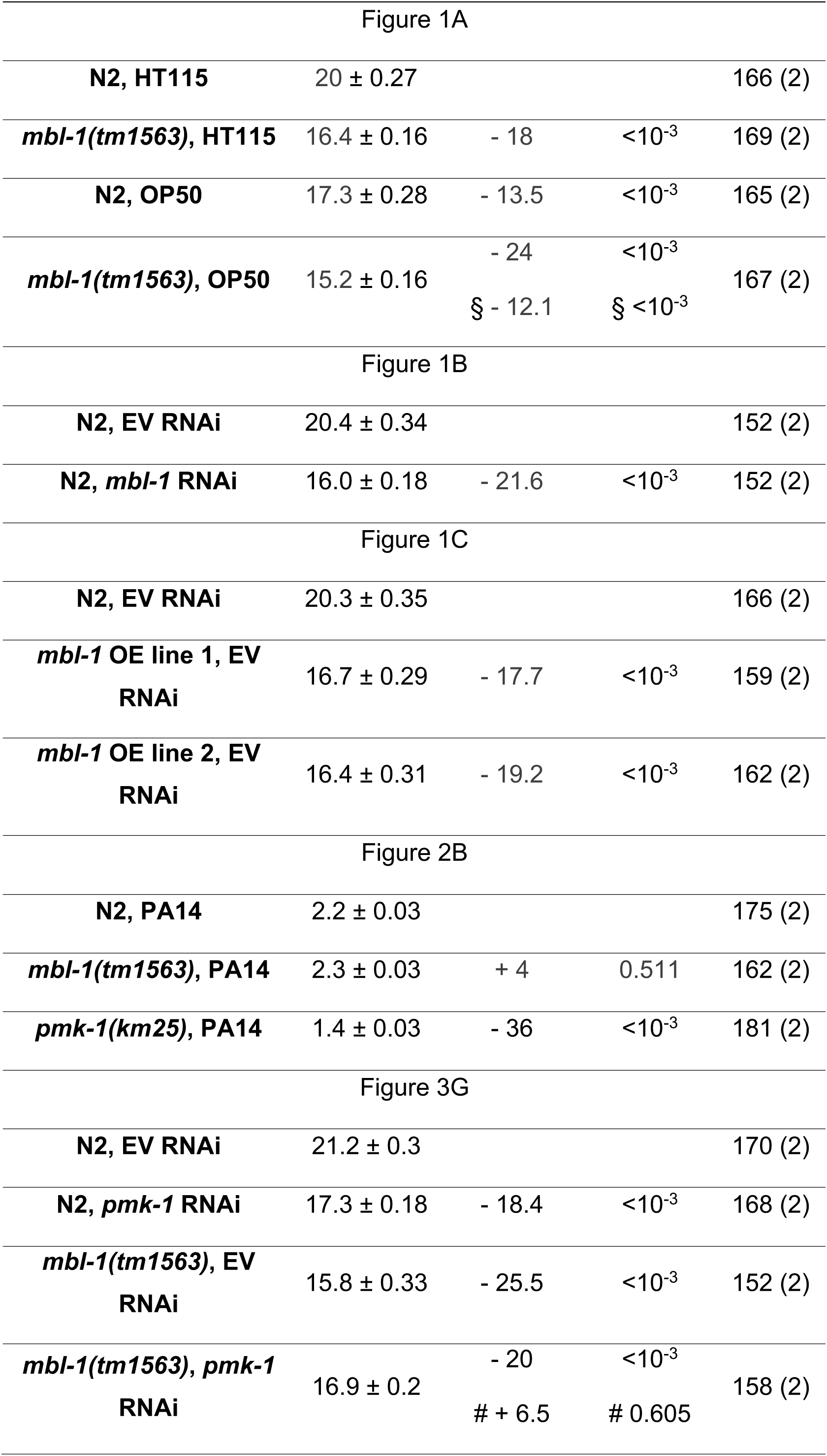
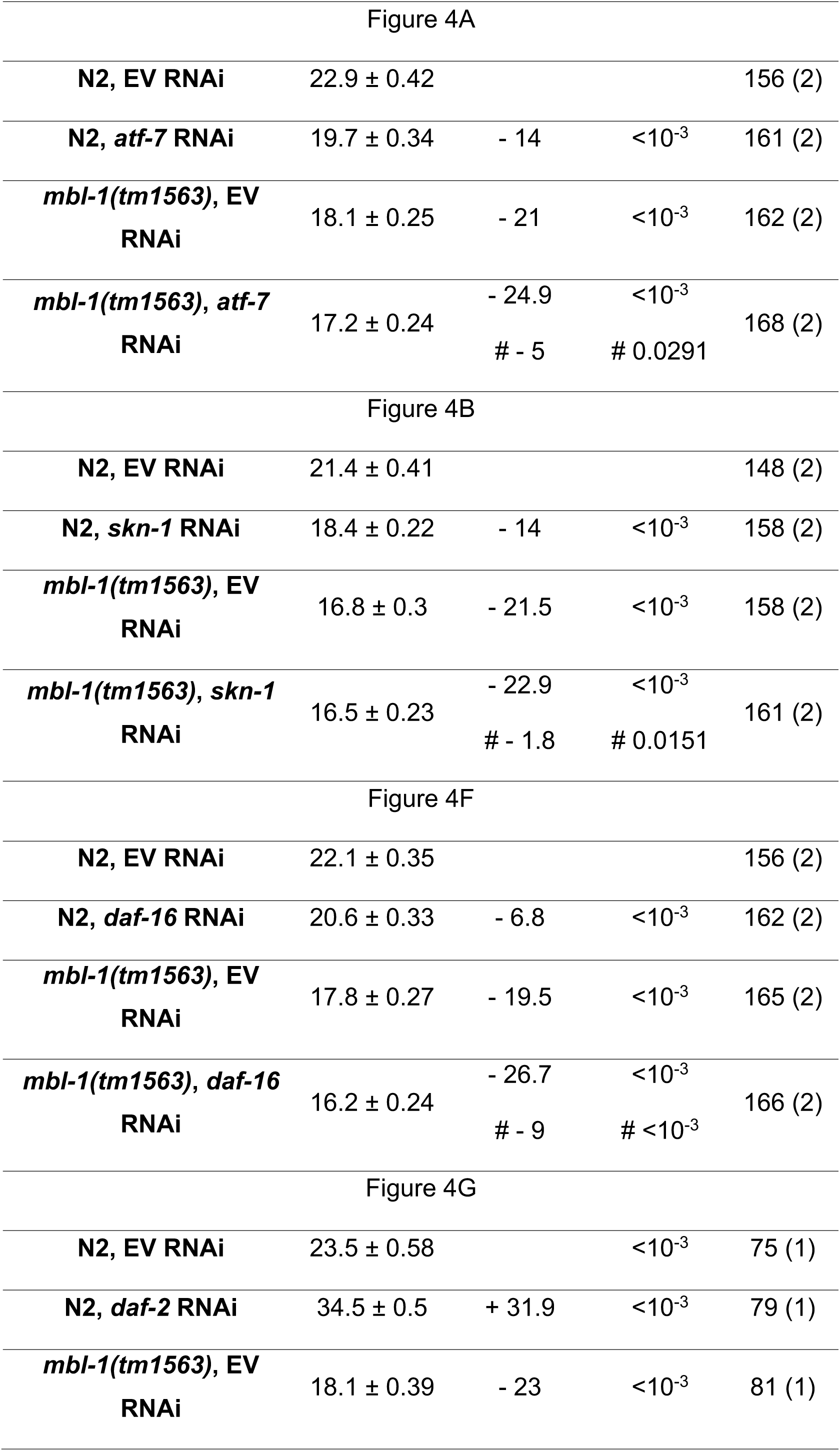
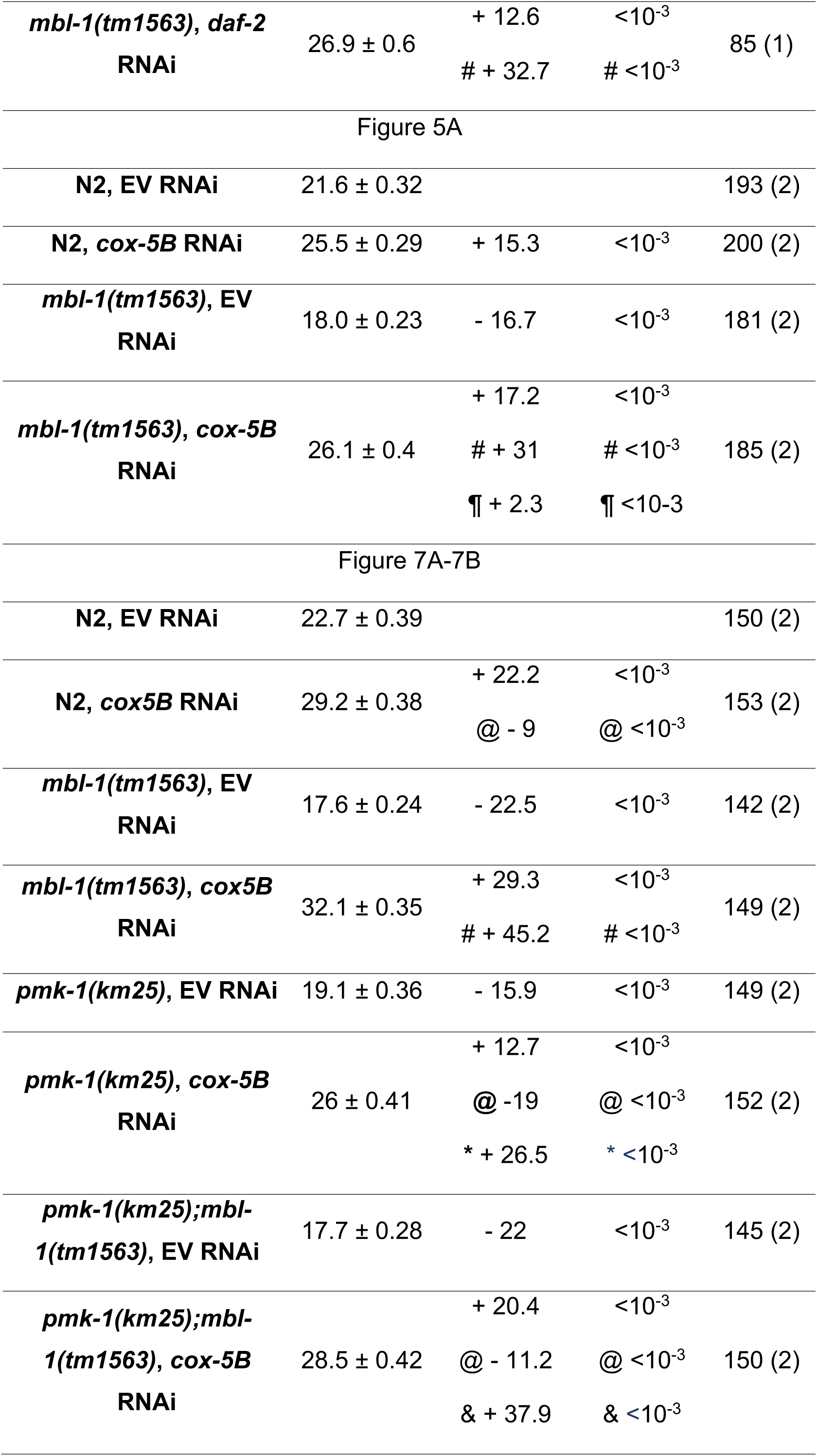
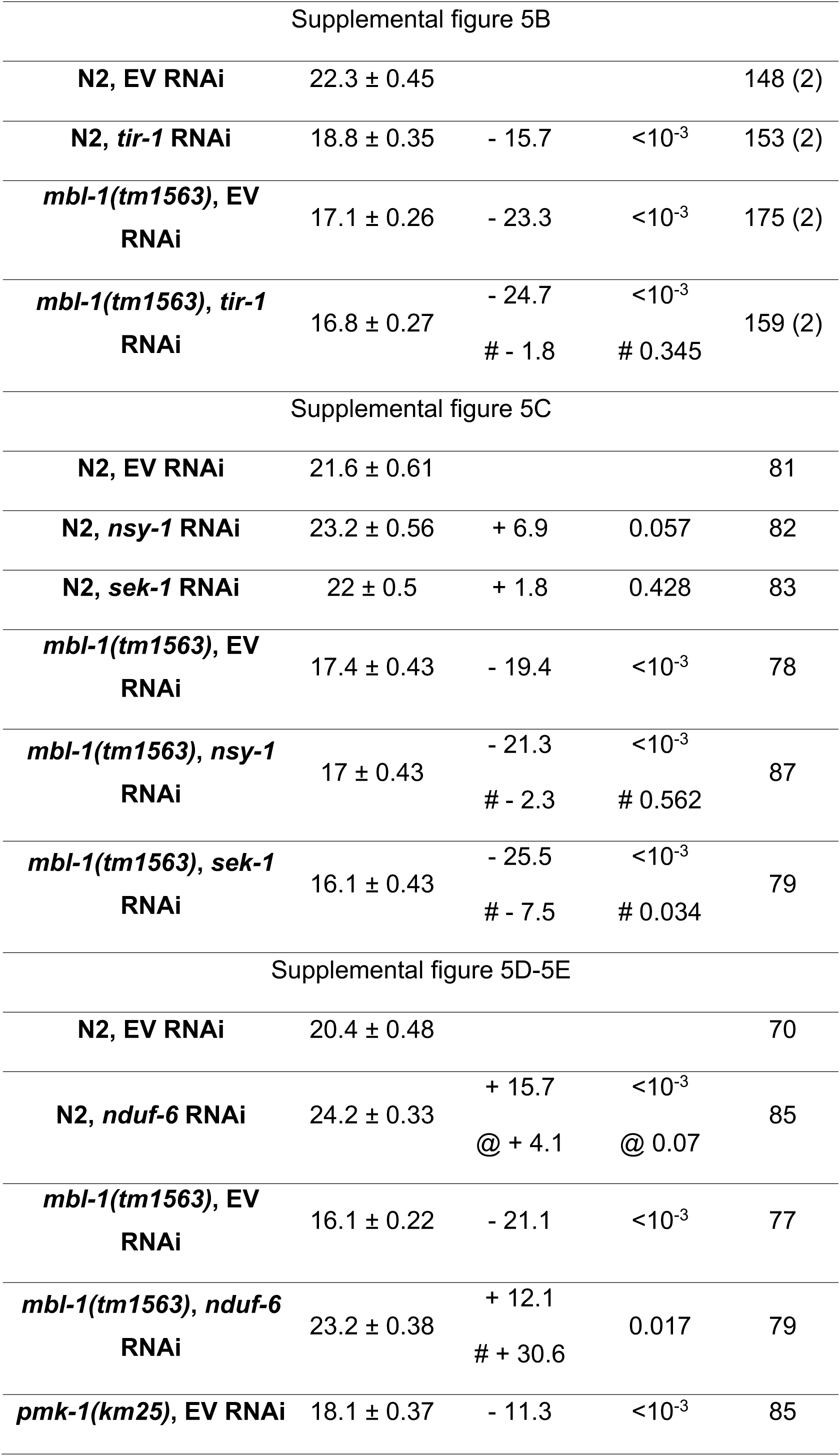
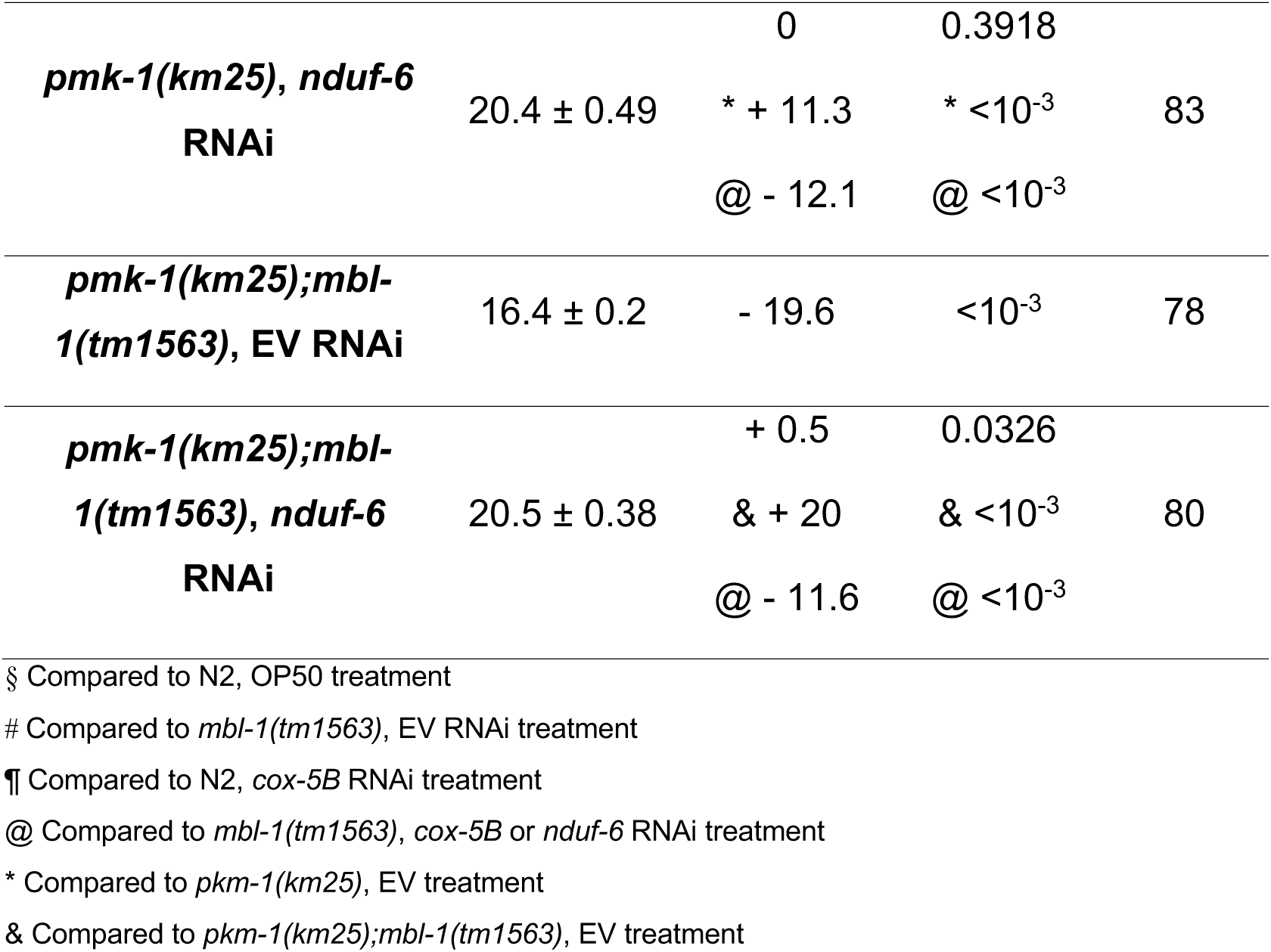
Summary of *C. elegans* lifespan experiments.

**S2 Table.**
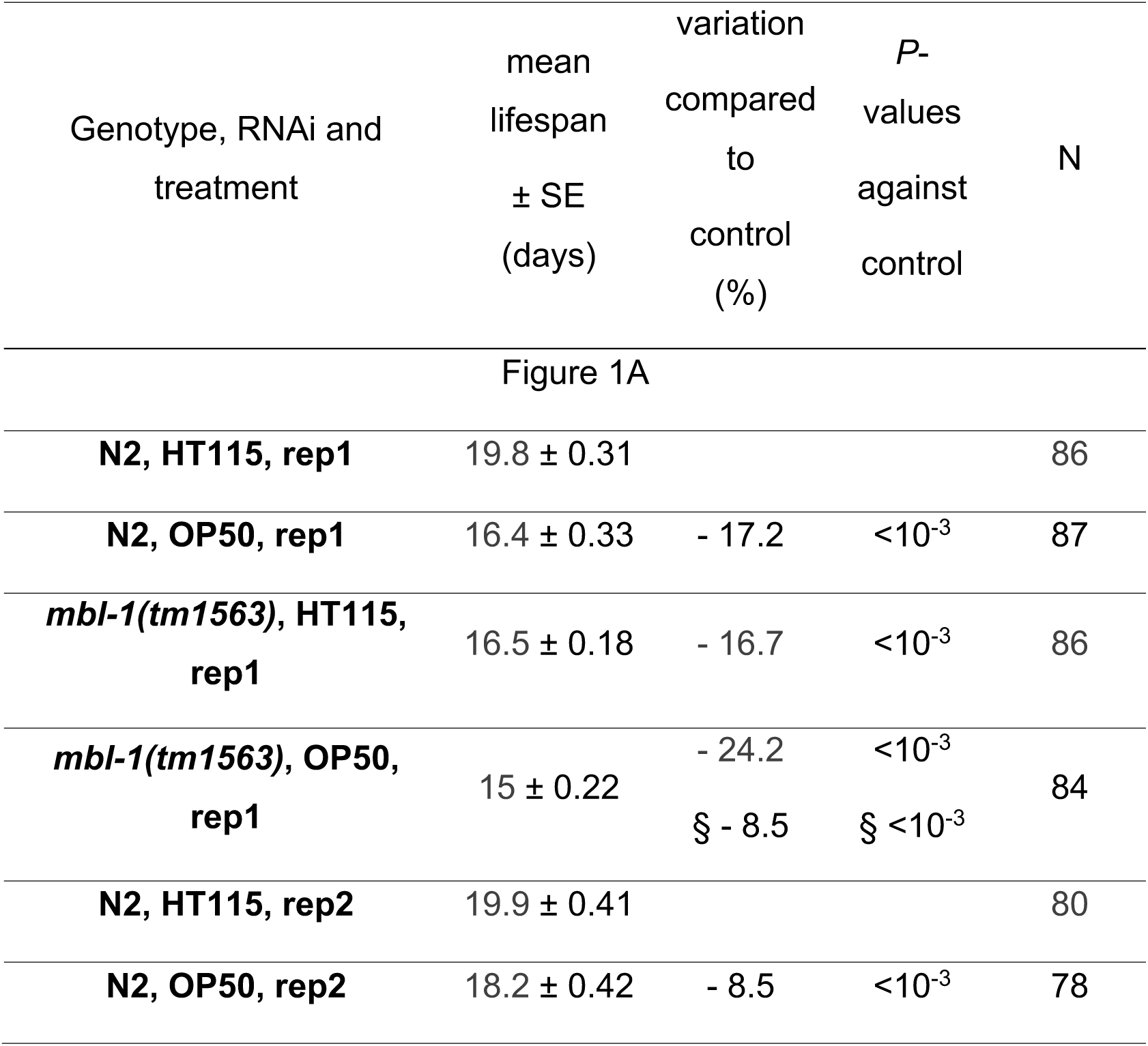
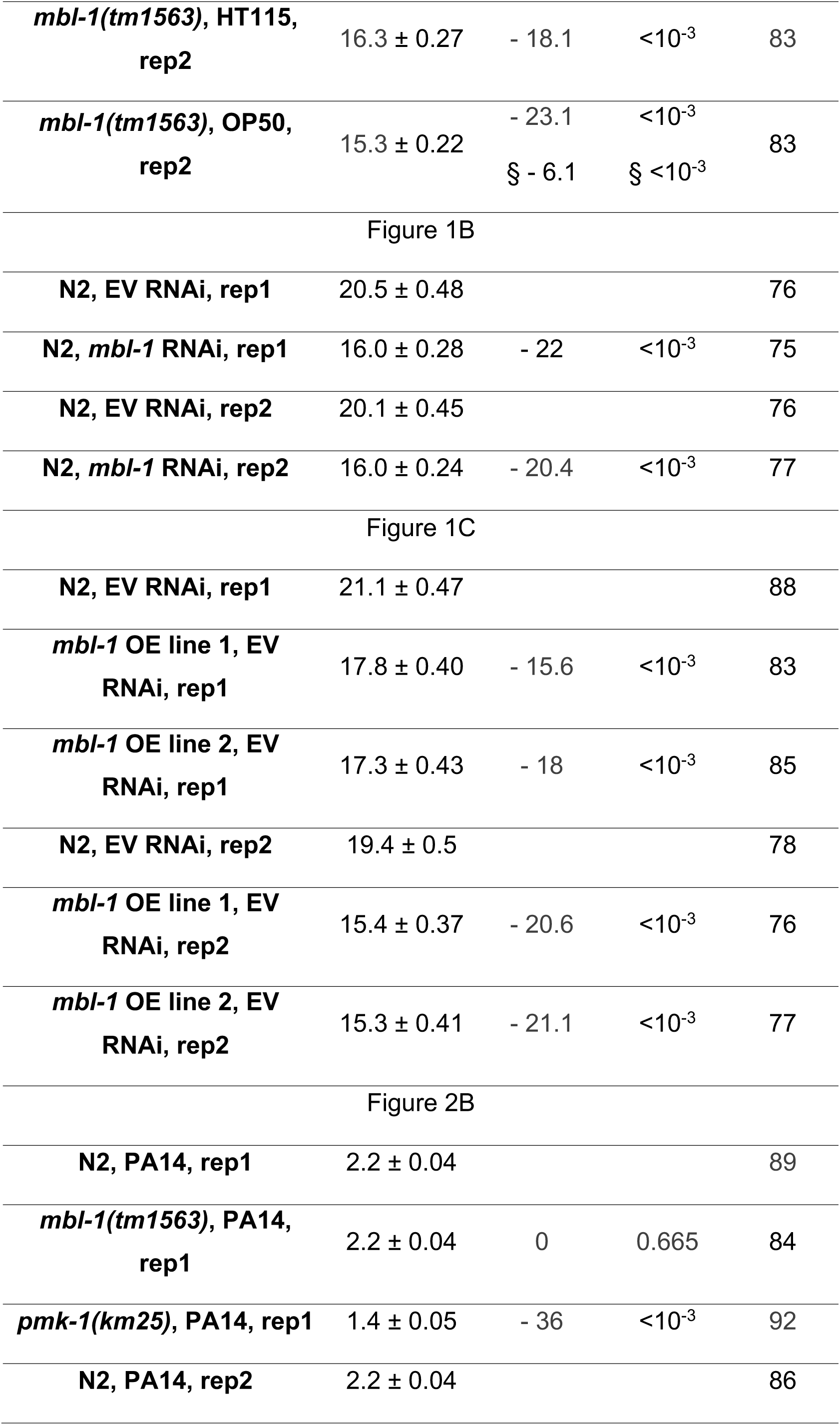
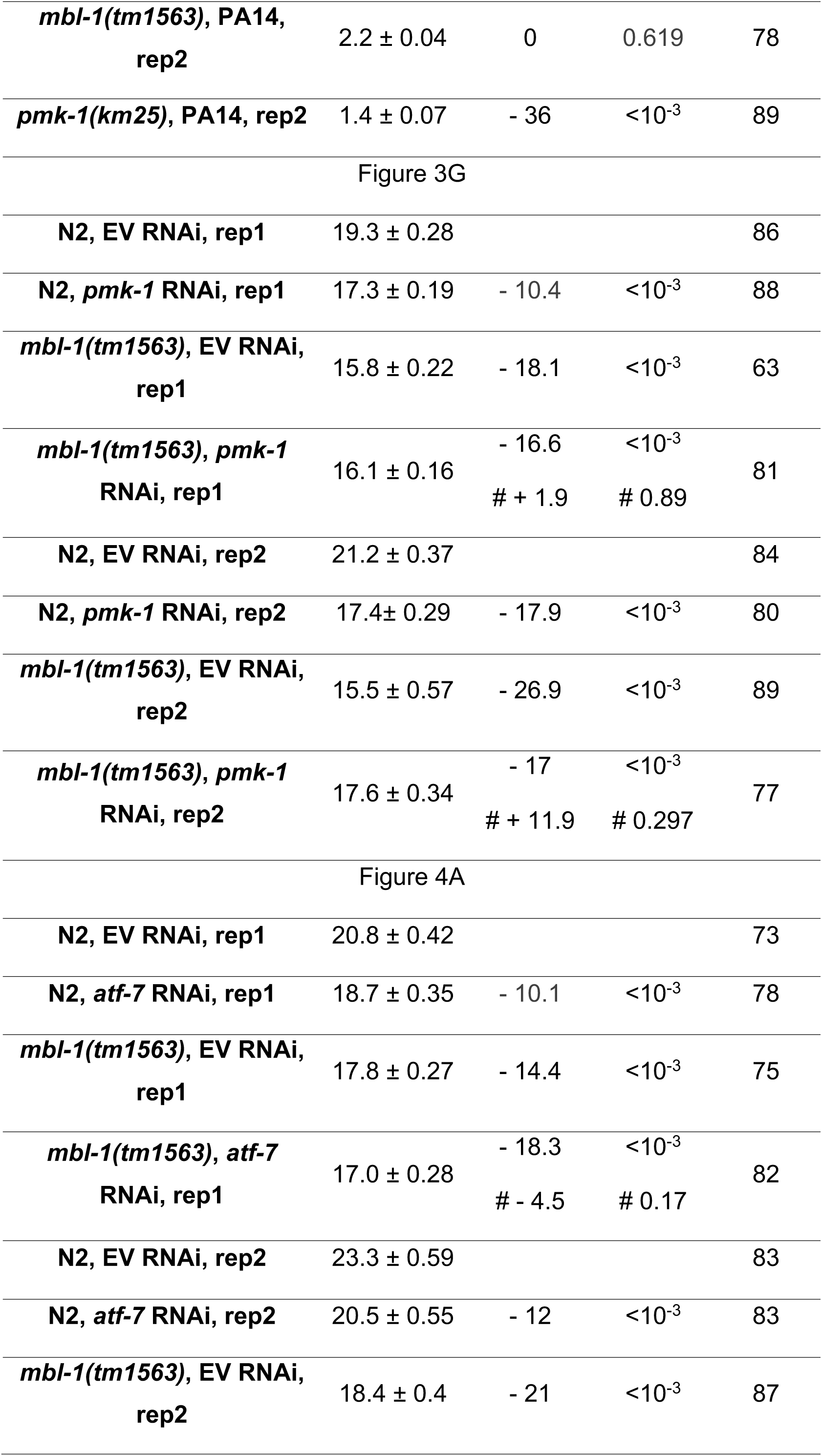
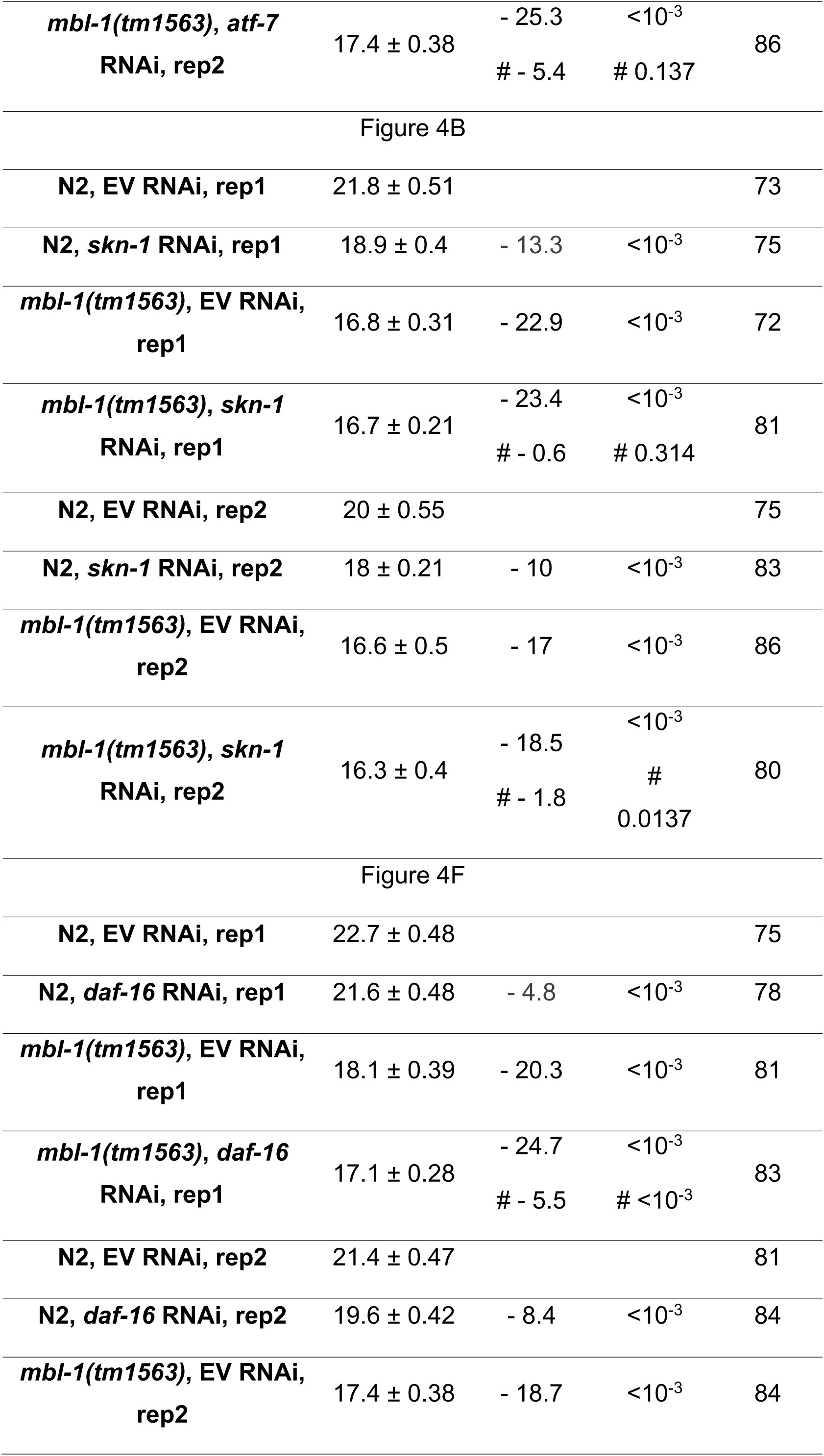
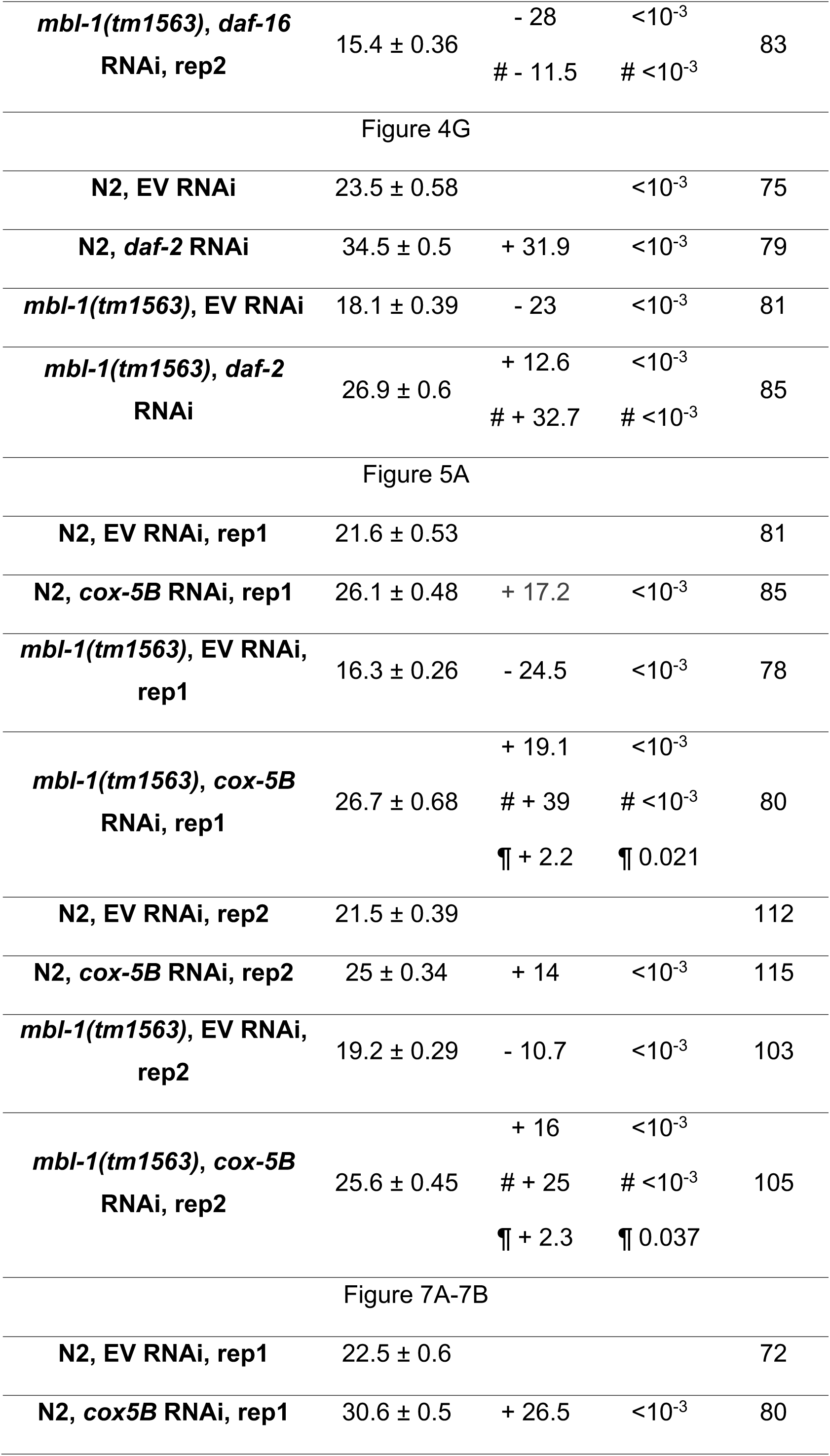
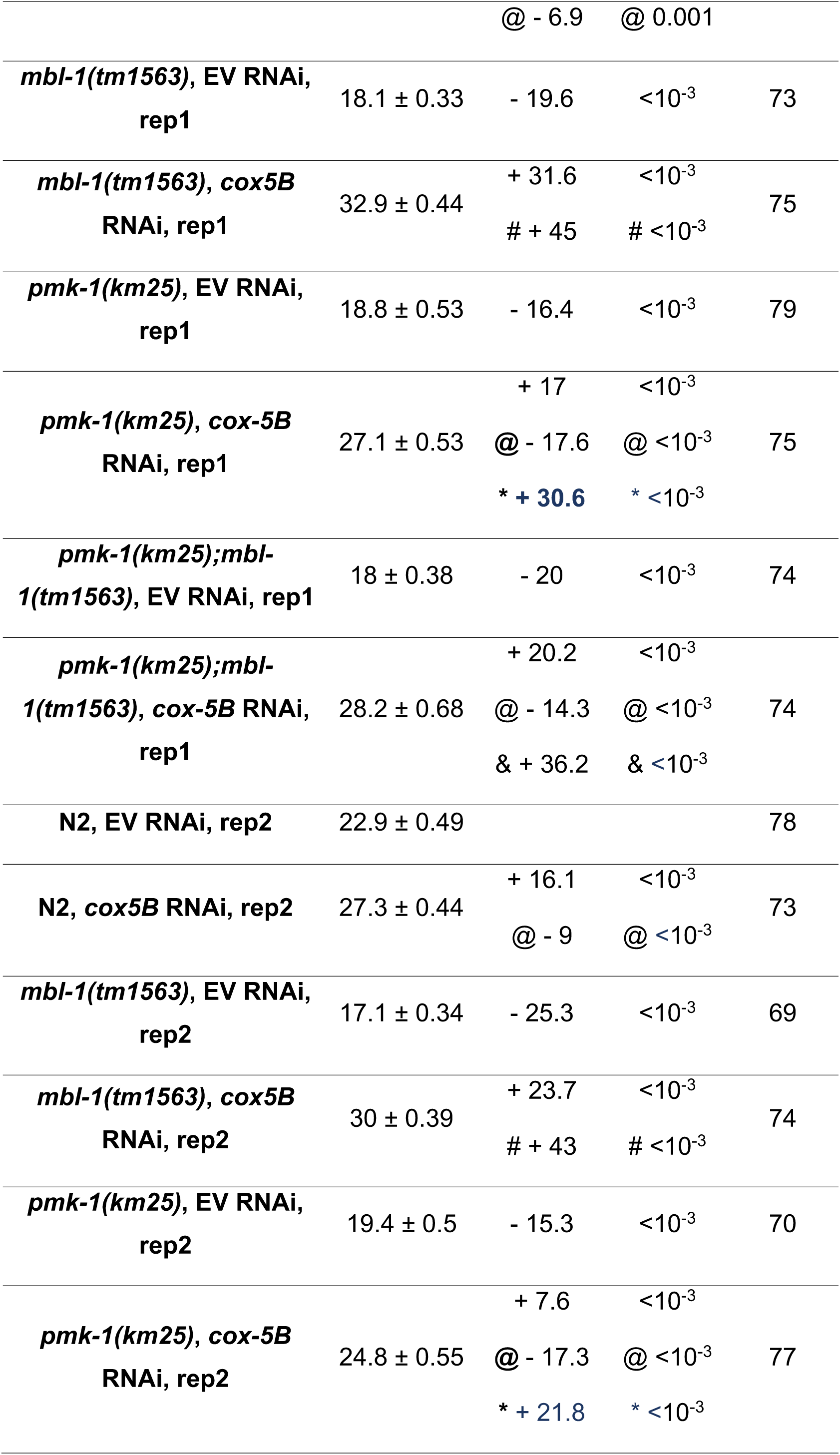
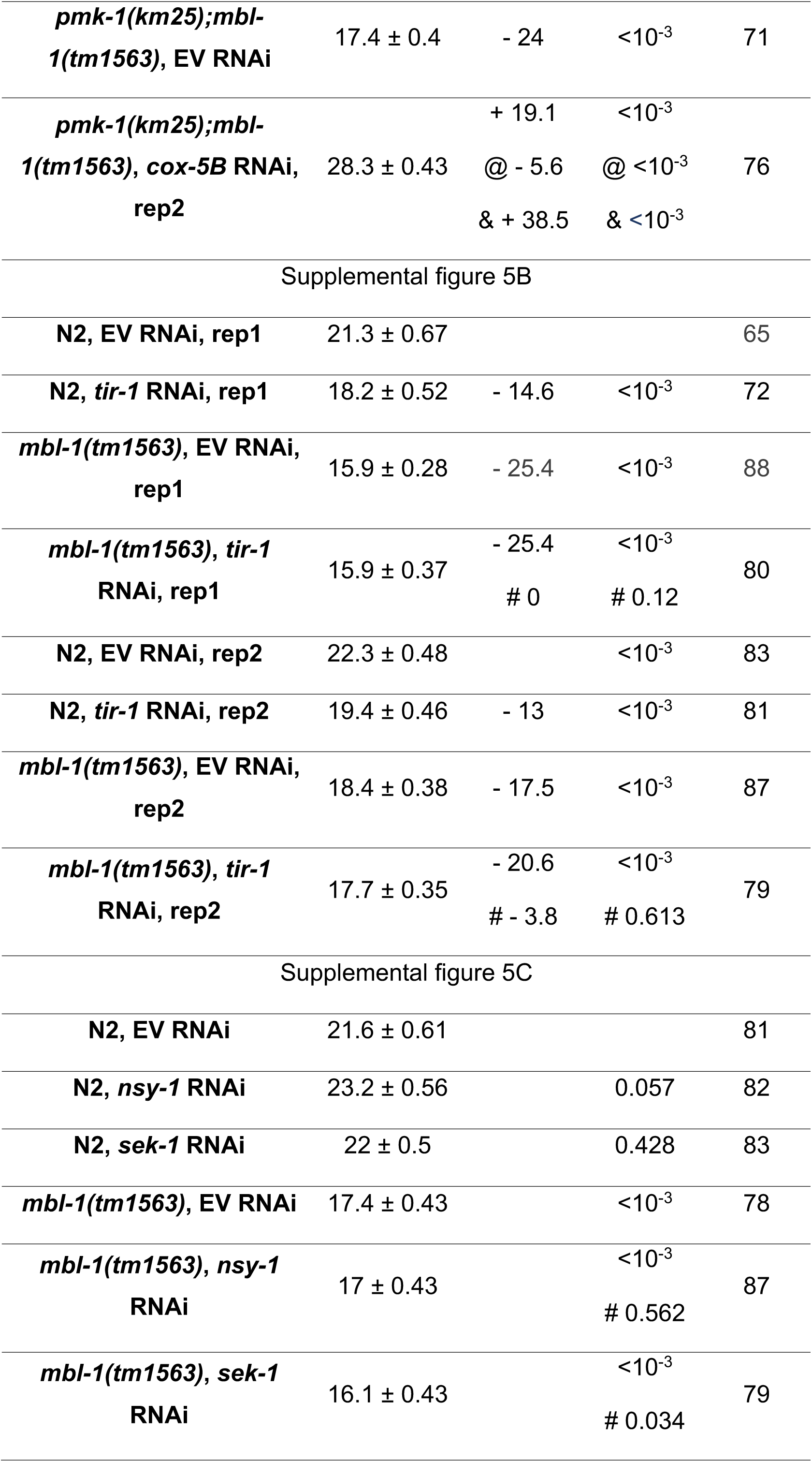
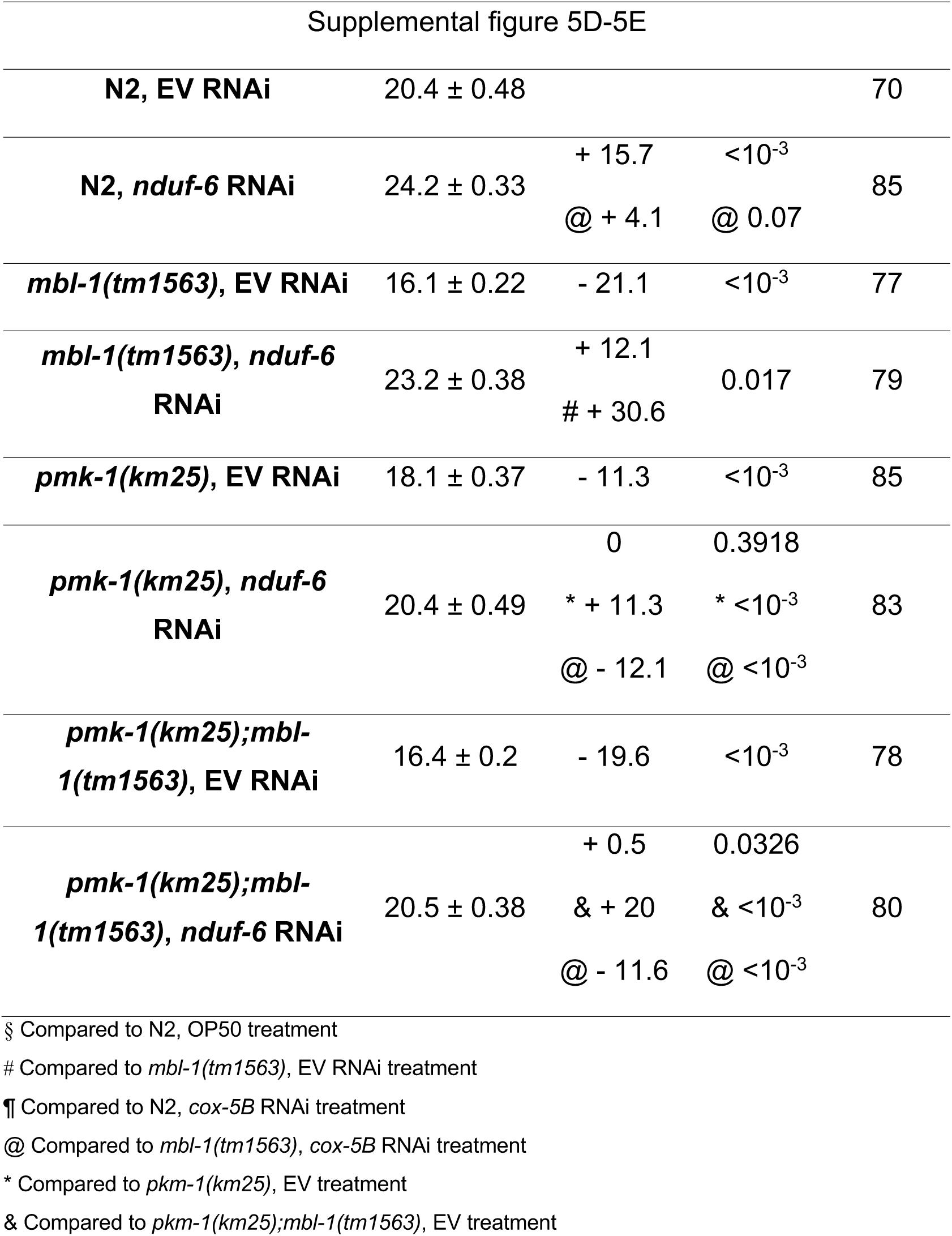
Individual replicates of *C. elegans* lifespan experiments.

**S3 Table.** Agarose gel quantifications. Supplemental Excel file.

**S4 Table.** List of up- and downregulated genes in day 2 adult *mbl-1(tm1563)* mutants. Supplemental Excel file.

**S5 Table.** WormExp output for up- and downregulated genes in day 2 adult *mbl-1(tm1563)* mutants. Supplemental Excel file.

**S6 Table.** Western blot quantifications. Supplemental Excel file.

**S7 Table.** Raw qRT-PCR and *gst-4p::gfp* imaging data. Supplemental Excel file.

**S8 Table.**
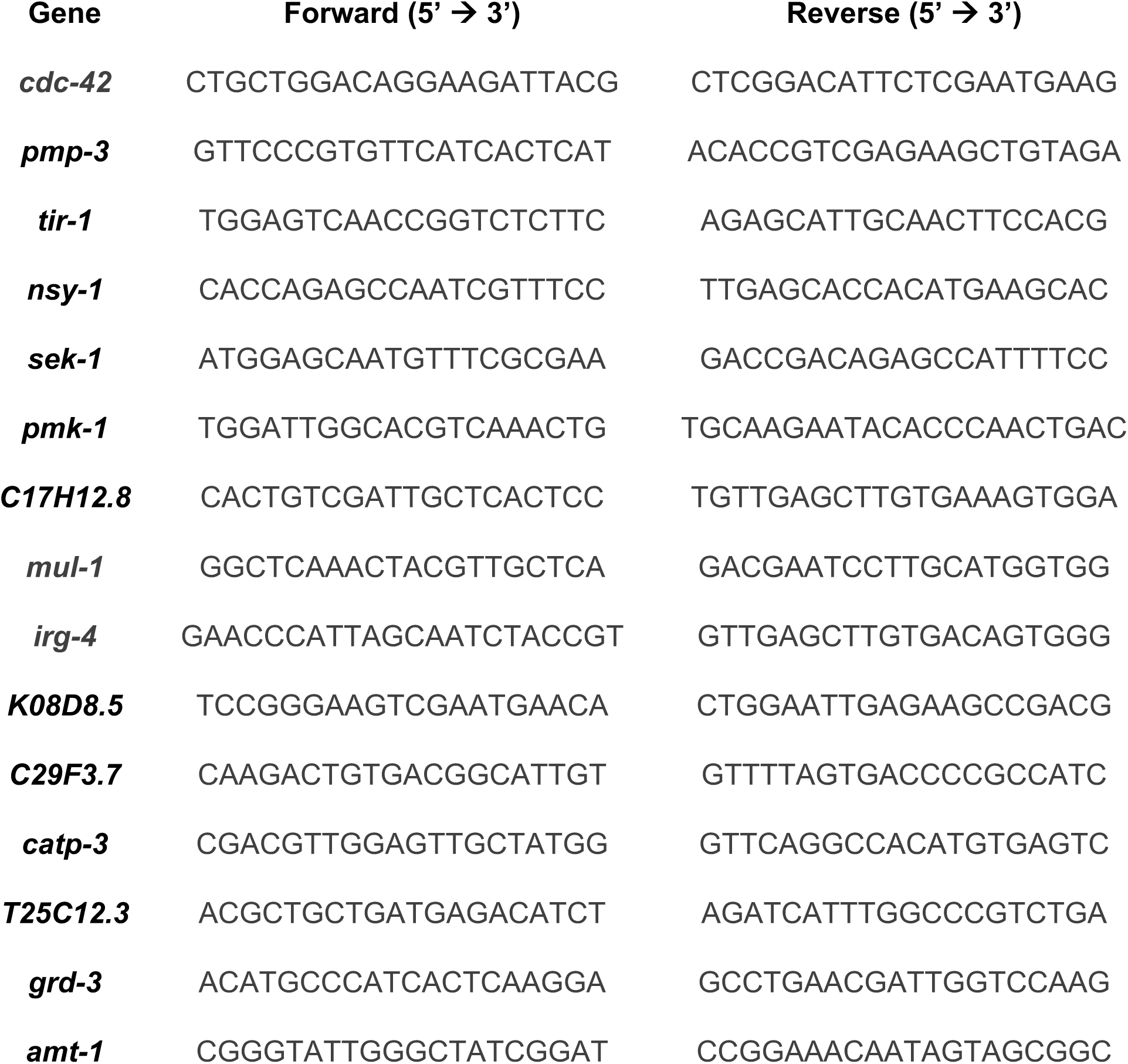
Oligonucleotide sequences used for qRT-PCR in this study.

**S9 Table.**
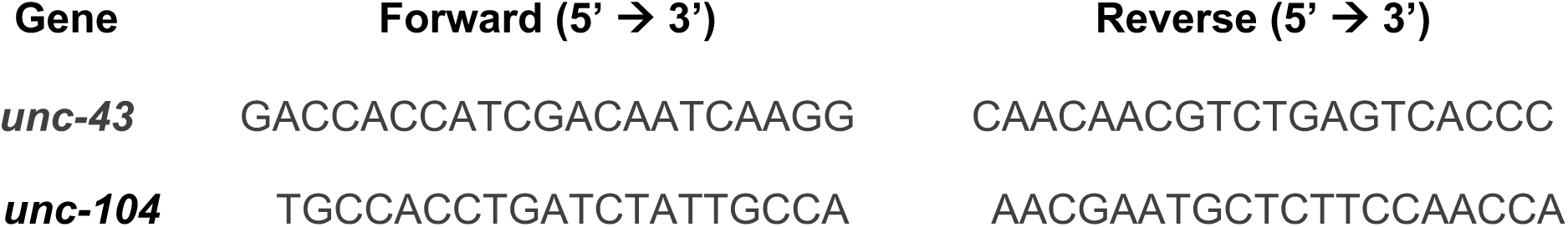
Oligonucleotide sequences used for splicing assay in this study.

